# Molecular evolution of a reproductive barrier in maize and related species

**DOI:** 10.1101/2024.12.02.626474

**Authors:** Elli Cryan, Garnet Phinney, Arun S. Seetharam, Matthew M.S. Evans, Elizabeth A. Kellogg, Junpeng Zhan, Blake C. Meyers, Daniel J. Kliebenstein, Jeffrey Ross-Ibarra

## Abstract

Three cross-incompatibility loci each control a distinct reproductive barrier in both domesticated maize (*Zea mays* ssp. *mays*) and its wild teosinte relatives. These three loci, *Teosinte crossing barrier1* (*Tcb1*), *Gametophytic factor1* (*Ga1*), and *Ga2*, each play a key role in preventing hybridization between incompatible populations and are proposed to maintain the barrier between domesticated and wild subspecies. Each locus encodes both a silk-active and a matching pollen-active pectin methylesterase (PMEs). To investigate the diversity and molecular evolution of these gametophytic factor loci, we identified existing and improved models of the responsible genes in a new genome assembly of maize line P8860 that contains active versions of all three loci. We then examined fifty-two assembled genomes from seventeen species to classify haplotype diversity and identify sites under diversifying selection during the evolution of these genes. We show that *Ga2*, the oldest of these three loci, was duplicated to form *Ga1* at least 12 million years ago. *Tcb1*, the youngest locus, arose as a duplicate of *Ga1* before or around the time of diversification of the *Zea* genus. We find evidence of positive selection during evolution of the functional genes at an active site in the pollen-expressed PME and predicted surface sites in both the silk- and pollen-expressed PMEs. The most common allele at the *Ga1* locus is a conserved *ga1* allele (*ga1-Off*), which is specific haplotype containing three full-length PME gene copies, all of which are non-coding due to conserved stop codons and are between 610 thousand and 1.5 million years old. We show that the *ga1-Off* allele is associated with and likely generates 24-nt siRNAs in developing pollen-producing tissue, and these siRNAs map to functional *Ga1* alleles. In previously-published crosses, the *ga1-Off* allele was associated with reduced function of the typically dominant functional alleles for the Ga1 and Tcb1 barriers. Taken together, this seems to be an example of a type of epigenetic trans-homolog silencing known as paramutation functioning at a locus controlling a reproductive barrier.

## Introduction

Reproductive barriers restrict gene flow between populations, facilitating both neutral and adaptive divergence. When populations diverge, these barriers can lead to speciation (Kulmuni et al. 2020). Reproductive barriers can be classified into either prezygotic barriers, which function before fertilization, or postzygotic barriers, which function after fertilization.

Prezygotic barriers are thought to generally be more complete, and thus more likely to lead to speciation, than postzygotic barriers (Christie et al. 2022; Baack et al. 2015). An important type of prezygotic reproductive barrier in plants is driven by post-pollination / pre-fertilization interactions between the pollen, the male gametophyte, and the pistil, the female floral structure. Pollen-pistil interactions can reduce gene flow between populations in a variety of flowering plants, including maize (Wang and Filatov 2023; Broz and Bedinger 2021).

Indigenous peoples domesticated maize (*Zea mays* ssp. *mays*) over nine thousand years ago in the Balsas valley region of Mexico (Piperno et al. 2009). At least two wild subspecies, the teosintes ssp. *Z. mays parviglumis* and *Z. mays mexicana*, played key roles in the origins of modern maize (Matsuoka et al. 2002; Yang et al. 2023). Farmers in Central America still cultivate maize alongside both of these taxa and other wild teosintes (Wilkes 1977). In some maize and wild teosinte populations, a group of three relatively common reproductive barriers controlled by the Gametophytic factor (GA) loci – *Tcb1*, *Ga1*, and *Ga2* – prevent gene flow between populations in only one direction to produce unilateral cross-incompatibility between populations (Kermicle 2006; Kermicle et al. 2006; Kermicle and Evans 2010).

The first of the three GA loci was characterized starting in 1901 when geneticists recorded phenotypic evidence, in the form of transmission distortion of the ratio of recessive sugary kernels, of a cross-incompatibility system in maize genetically controlled by a locus now known as *Gametophytic factor1* (*Ga1*) (Corrins 1901; Mangelsdorf and Jones 1926; Schwartz 1950). The *Ga1* locus encodes two tightly linked gametophytic factors (genes) whose products interact after pollination but before fertilization. One gene generates an active prezygotic reproductive barrier in the female floral organ, the silk, and a matching second gene enables the male gametophyte, the pollen, to overcome that barrier [Figure 1]. The silk and pollen-expressed genes each encode distantly related pectin methylesterases (PMEs). PMEs play important roles in plant cell growth by enzymatically modifying cell wall pectin properties, impacting cell wall growth dynamics, especially in rapidly-growing plant cells like those in both the pollen tube and silk tissues (Shin et al. 2021; Wallace and Williams 2017). When both the silk and pollen *Ga1* genes are active, the *Ga1* pollen tube can grow normally down the transmitting tract in the *Ga1* silk to eventually reach the female gametophyte, and fertilization can occur. In contrast, when the *Ga1* silk gene is active but the pollen gene is inactive, the *Ga1* silk impedes *ga1* pollen tube growth, possibly through the PME altering the integrity of the pollen tube cell wall and inhibiting directional growth. This inhibition reduces the chances of or prevents fertilization, producing the Ga1 reproductive barrier. The barrier only prevents gene flow from *ga1* to *Ga1* plants; in the opposite direction, GA active pollen can grow normally, although more slowly than GA inactive pollen, down a GA inactive silk (Lu et al. 2014). Study of *Ga1* is complicated by the fact that the locus contains a complex and polymorphic pattern of duplicated pseudogenes (Bapat et al. 2023).

**Figure 1:**
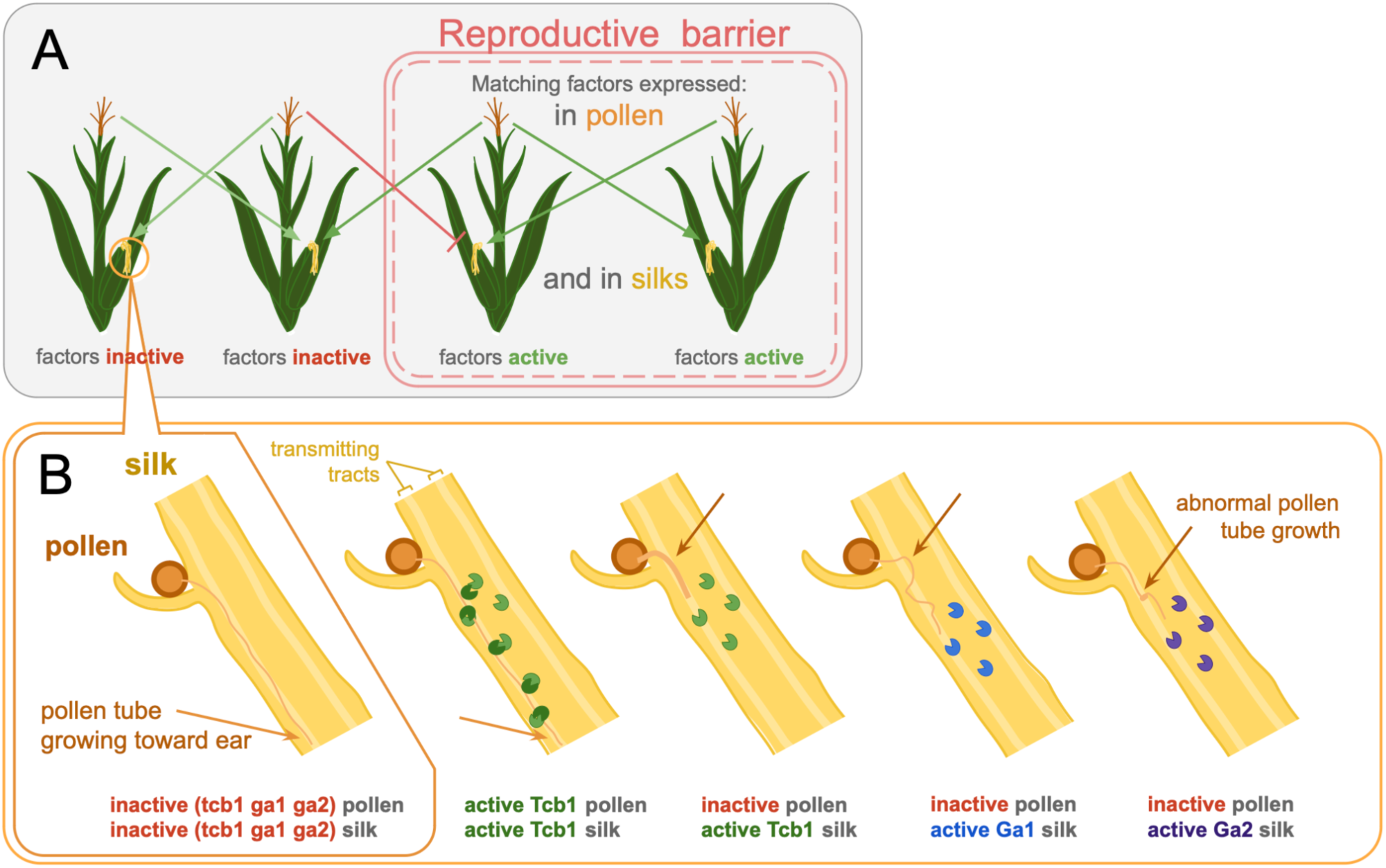
Incompatibility between GA inactive pollen and GA active silk can generate a reproductive barrier between *Zea mays* populations. (A) When a gametophytic factor (GA) gene is active in the silk, only pollen with a matching active GA gene can grow normally down the silk toward the female gametophyte (light green arrow), which impedes the chances of fertilization by inactive GA pollen (dark red line) and generates a reproductive barrier. (B) Diagram based in part on microscopy published by Lu, Kermicle, and Evans 2014: When silk and pollen both have no GA gene activity or both have matching active GAs, the pollen tube grows quickly down a transmitting tract in the silk toward the ear. Each of the three silk GAs impacts inactive GA pollen tube morphology differently. Tcb1 silk PMEs are shown in green, Ga1 in blue, and Ga2 in purple. Tcb1 pollen PMEs, shown in dark green, indirectly or directly interact with silk PMEs.

Following the characterization of the Ga1 barrier and locus, maize geneticists identified and validated two additional GA loci, named *Teosinte crossing barrier1* (*Tcb1*) and *Ga2* (Evans and Kermicle 2001; Brieger 1937; Burnham 1936). Each locus functions similarly to *Ga1*, encoding a silk PME gene and a tightly linked, distantly related, matching pollen PME gene. For clarity, we call the genes active in the silk *Tcb1k, Ga1k,* and *Ga2k*, and the genes active in the pollen *Tcb1p*, *Ga1p*, and *Ga2p* [Table 1]. In spite of the genetic and mechanistic similarity between loci, the three loci are distinct in the sense that each silk gene-encoded GA barrier can only be fully overcome by pollen with a matching active GA pollen gene, although pollen with a mismatched active GA gene is slightly preferred to pollen with no active GA genes (Lu et al. 2019). Additionally, the morphology of an inhibited wildtype pollen tube differs depending on whether the *Tcbk*, *Ga1k*, or *Ga2k* gene is active in the silk, suggesting that each silk gene may have a slightly different molecular function [Figure 1] (Lu et al. 2014).

**Table 1:** GA locus, allele, and gene nomenclature. Each GA barrier is controlled by a locus containing corresponding silk- and pollen-expressed genes. GA alleles can be categorized by activity of the PMEs, or factors, encoded by the genes in each locus. Here, we propose a new *ga1-O* allele, which is distinct from a fully inactive ga1 allele. Weakly active Tcb1, Ga1, and Ga2 barriers have been observed, but often under the control of alleles which could be called strong in other genetic backgrounds. Alleles like these have sometimes been called *Ga1-W* and *Ga2-W*; to date, no *Tcb1-W* allele has been characterized.

| Tcb1 |  |  | Ga1 |  |  | Ga2 |  |  |
| --- | --- | --- | --- | --- | --- | --- | --- | --- |
| Alleles | *Tcb1k* (silk) activity | *Tcb1p* (pollen) activity | Alleles | *Ga1k* (silk) activity | *Ga1p* (pollen) activity | Alleles | *Ga2k* (silk) activity | *Ga2p* (pollen) activity |
| *Tcb1-S (Strong)* | active | active | *Ga1-S (Strong)* | active | active | *Ga2-S (Strong)* | active | active |
| *Tcb1-W (Weak)* | weakly active | active | *Ga1-W (Weak)* | weakly active | active | *Ga2-W (Weak)* | weakly active | active |
| *Tcb1-M (Male)* | inactive | active | *Ga1-M (Male)* | inactive | active | *Ga2-M (Male)* | inactive | active |
| *tcb1* | inactive | inactive | *ga1-N (Null)* | inactive | inactive | *ga2* | inactive | inactive |
|  |  |  | *ga1-O (Off)* | inactive *Ga1k*-targeting siRNA | inactive |  |  |  |

Although the GA loci have been studied for generations, the repetitive complexity of the loci and resulting recalcitrance to sequencing have long impeded research on the molecular evolution of the GA barriers. In the absence of high-quality and complete gene sequence data, ecological and modeling research suggested different evolutionary histories for the GA loci.

Because the barriers are commonly observed in sympatric teosinte populations, many authors have argued that the GA loci evolved to keep maize and teosinte distinct (Evans and Kermicle 2001; Zhang et al. 2023). However, population dynamics modeling work suggests that GA-like loci are largely unable to sustain a long-term crossing barrier between populations in an annual plant (Rushworth et al. 2022). Recent advances in sequencing have enabled the identification of all six types of genes controlling the gametophytic factors in maize, such that one reference gene sequence exists each for *Tcb1k*, *Tcb1p*, *Ga1k*, *Ga1p*, *Ga2k*, and *Ga2p* (Lu et al. 2019; Zhang et al. 2023; Moran Lauter et al. 2017; Zhang et al. 2018; Chen et al. 2022). Simple sequence comparisons suggest the origins of the *Ga1k* and *Tcb1k* genes long predate the domestication of maize ∼9k years ago (Bapat et al. 2023; Zhang et al. 2023) The evolution of GA loci is complicated further by evidence of the attenuation of incompatibility in some backgrounds (Lu et al. 2014; Lu et al. 2019; Goodman et al. 2021; Demerec 1929; Nelson 1952; Ashman 1975) and the observation of a complex series of highly repetitive haplotypes at the *Ga1* locus (Bapat et al. 2023).

Combined, these data motivate our detailed assessment of the diversity, function, and evolution of all the GA loci and how they may have impacted the evolution of *Zea*. Here, we work toward this goal by analyzing all three loci using long-read genome assemblies to improve reference gene models, identifying GA genes in more than fourteen new species, and classifying the diversity of GA genes and loci. We establish a better understanding of the evolution and function of the GA system by estimating the timing of the evolution of the loci, testing for selection on genes and loci, and documenting the location of selected sites on predicted protein structure. Surprisingly, we find evidence for epigenetic silencing of the GA loci associated with an inactive *ga1* allele we call *ga1-Off*.

## Results

### Refined models of gametophytic factor (GA) genes

Though individual reference gene sequences have been published for both silk and pollen PME genes for all three gametophytic factor loci, the full genomic regions for all three loci are largely unannotated across available genomes (Lu et al. 2019; Zhang et al. 2023; Moran Lauter et al. 2017; Zhang et al. 2018; Chen et al. 2022). To study the full sequences of all three GA loci, we identified their genomic regions in 52 genomes spanning *Zea* and related genera [Figure 2, Supplemental table 1]. Our initial genome-wide BLAST search for the reference GA gene sequences did not find the published *Ga2k* and *Tcb1p* reference genes in genomes of plants known to produce the Ga2 barrier and overcome the Tcb1 barrier, functions encoded respectively by *Ga2k* and *Tcb1p*. We shifted to a synteny-based approach, using known genes from loci flanking each GA locus in the maize genomic map to identify the genomic coordinates of a large region that should contain the full GA locus. Using this approach, we were able to identify improved reference gene sequences for *Ga2k* and *Tcb1p.* These sequences share features of the other GA genes – including intron exon structure, expression patterns, signal peptide presence, and amino acid similarity – and they are present and expressed in reproductive tissues of plants with barrier function (see methods). Our improved version of the *Ga2k* gene model is shorter than the previously verified *Ga2k* sequence sourced from a BAC, which includes 3’ sequence missing from genomes of maize plants that generate the Ga2 barrier [Supplemental figure 1] (Chen et al. 2022). Our *Tcb1p* gene model has a shifted intron position, which has little to no impact on amino acid sequence, but better allowed us to identify *Tcb1* loci because the nucleotide sequence is more consistent across species [Supplemental figure 1] (Zhang et al. 2023).

**Figure 2:**
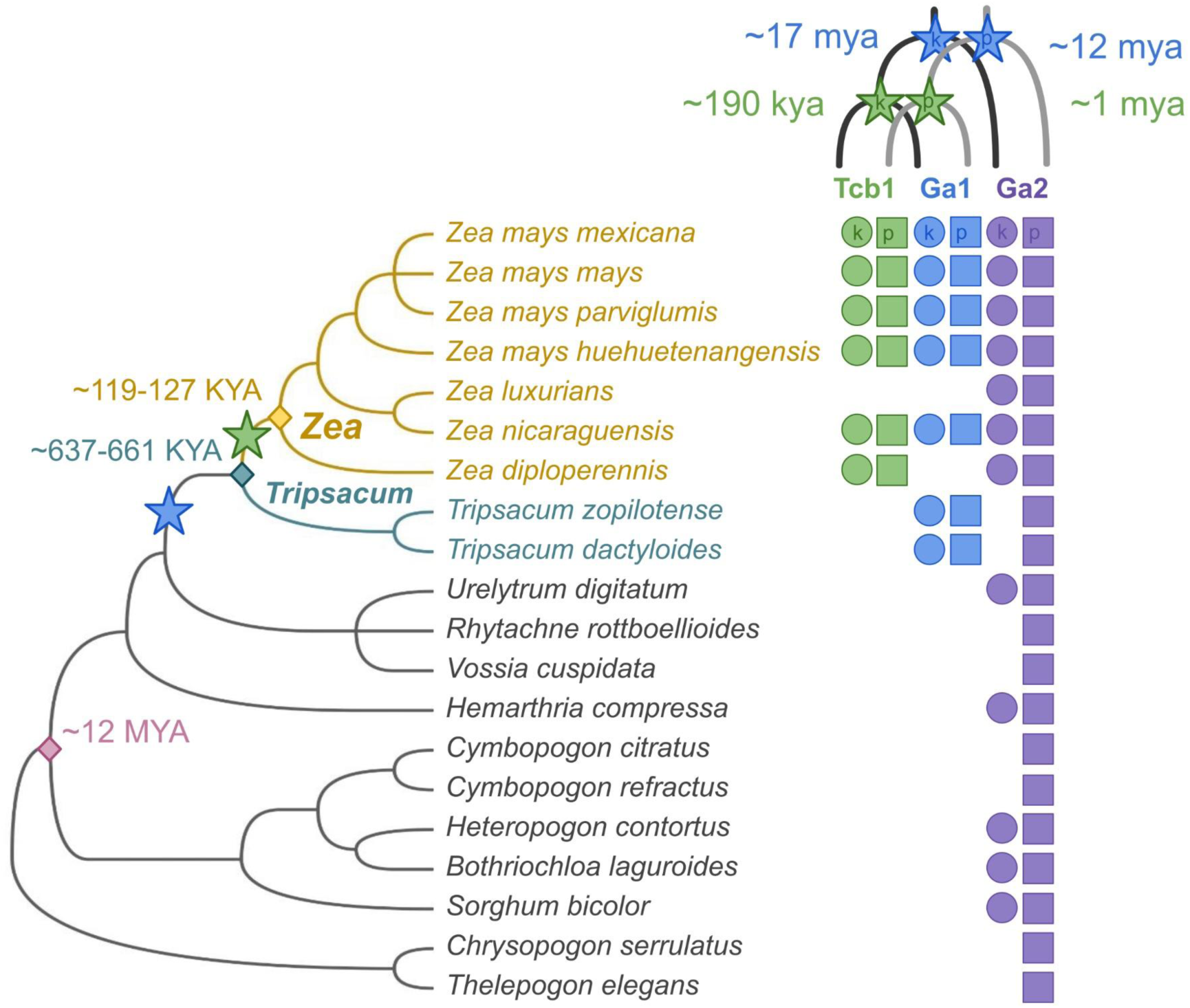
GA gene homologs are present in domesticated maize, three teosinte subspecies, and sixteen other related grasses. Presence of GA genes is indicated by circles for silk-expressed (k) and squares for pollen-expressed (p) genes. Species divergence times are from (Chen et al. 2022; Welker et al. 2020), and the species tree is from (Grass Phylogeny Working Group III 2024). Divergence of *Ga1/Tcb1* from *Ga2* is indicated by a blue star and divergence of *Tcb1* from *Ga1* is indicated by a green star. Gene divergence times are based on sequence dissimilarity and Ks, see Supplemental table 2.

Our confidence in our improved gene models, and in our understanding of all three GA loci, comes in part from P8860, an inbred maize line with functional barrier loci (*Ga1-M*, *Tcb1-S*, *Ga2-S*) introgressed from wild relatives (*Ga1-M* and *Tcb1-S* are from Collection 48703 of *Zea mays mexicana* (Kermicle and Allen 1990) and *Ga2-S* is from plant 3 in Collection 104 of *Zea mays parviglumis* (Kermicle and Evans 2010). We assembled P8860 using long read sequencing (see methods for assembly details). Our improved *Ga2k* and *Tcb1p* gene models, as well as previously published reference sequences for the other active genes, match sequences present in this genome [Supplemental figure 2]. Many maize lines have no or only one active barrier, and this is the first reference-quality genome assembly of a maize line with the Tcb1 barrier active.

### GA genes are present in diverse maize and wild relative genomes

To date, all three GA barrier phenotypes have only been reliably documented in subspecies of *Zea mays*. Using our updated gene models, we identified GA loci in 29 high quality genome assemblies of maize lines and 23 genomes of related wild taxa from both *Zea* and eleven related *Andropogoneae* genera [Figure 2]. We focused interpretation on the presence of the locus in non-*Zea* taxa because absence of GA loci in these genomes reflect either true absence or false negatives arising from incomplete scaffolding. In general, these loci contain many truncated gene fragments and only a few full-length gene copies, and the existing full-length gene models within the loci were unannotated in many of the genomes. All three loci, when present, are syntenic to the corresponding locus in *Zea mays*, and when silk genes are present, a tightly linked pollen gene is also found within the locus. These efforts represent the first comprehensive identification of *Ga1* and *Ga2* in non-Zea species.

Mapping the presence of the GA loci onto a phylogeny of the *Andropogoneae* tribe (Welker et al. 2020; Grass Phylogeny Working Group III 2024) indicates that *Tcb1* likely arose after the divergence of *Zea* from *Tripsacum* approximately 650K years ago, but before or around the time of the diversification of taxa within *Zea* around 170K years ago [Figure 2] (Chen et al. 2022b). This clearly predates the divergence of the three wild subspecies of *Zea mays*, the teosinte ssp. *mexicana*, *parviglumis*, and *huehuetenangensis*, which first split 30-60K years ago (Chen et al. 2022b). Similar reasoning suggests that Ga1, present in *Zea* and *Tripsacum* but not in other genera, arose more than 650K years ago, but likely after the Andropogoneae tribe arose ∼14 [9.89-17.97] million (M) years ago (Chen et al. 2022; Welker et al. 2020). We also estimated gene divergence times as an independent way of dating the origin of each locus. We used pairwise alignment of gene model sequences to calculate a synonymous substitution rate and estimated time since divergence, assuming one generation per year and a standard maize mutation rate of μ = 3.3 * 10^-8 [Supplemental table 2] (Clark et al. 2005). The gene divergence times are mostly consistent with the locus divergence timing estimates from the species phylogeny [Supplemental figure 3, Supplemental table 2]. We estimate that *Ga1k* diverged from *Ga2k* around 17M years ago and *Ga1p* diverged from *Ga2p* around 12M years ago, supporting the idea that *Ga1* arose from a duplication of *Ga2* ∼12-17M years ago, consistent with previous work (Lu et al. 2019) [see Figure 2]. *Ga1* silk and pollen gene divergence times are similar, supporting the idea that silk and pollen genes were already tightly linked since before *Ga1* arose. In contrast, our estimates of the age of the *Tcb1* pollen and silk genes differ by an order of magnitude. While *Tcb1k* seems to have diverged from *Ga1k* ∼190K years, consistent with silk gene presence in the species phylogeny, our estimate of divergence time between *Tcb1p* and *Ga1p* is ∼1M years and predates the origin of the *Zea* genus. Although the timing of the *Ga2* duplication that led to *Ga1* is roughly concurrent with an ancient allopolyploidization event ∼10M years ago in the Tripsacinae lineage (Wang et al. 2015), *Ga1* and *Ga2* do not share synteny beyond the local boundaries of the GA loci, suggesting that the Tripsacinae whole genome duplication was not the source of *Ga1* [Supplemental figure 4].

### GA loci exhibit high haplotypic diversity within Zea

To assess the functional variation within and between each of the GA loci, we characterized the complete haplotypes of each GA locus in our sampled genomes by documenting the sequence similarity and gene order of full-length gene copies at each locus [Figure 3]. All three loci show variation in functional gene copy number, ranging from 0 to 2 for silk genes and 0 to 8 for pollen genes. While most haplotypes were found in a single individual, others were shared across up to eighteen genomes. Given the observation that there are individual genotypes with functional gene sequences but non-functional barriers, we combined our sequence analysis of genome assemblies with functional genomic data for methylation and expression available for many of these genomes to assess potential epigenetic differences. At all three loci, when the barrier is active the silk gene is unmethylated, and when the pollen can overcome the barrier the pollen genes are highly expressed, though sometimes display methylation in diploid tissue. Below, we discuss the specific observations for each locus individually.

**Figure 3:**
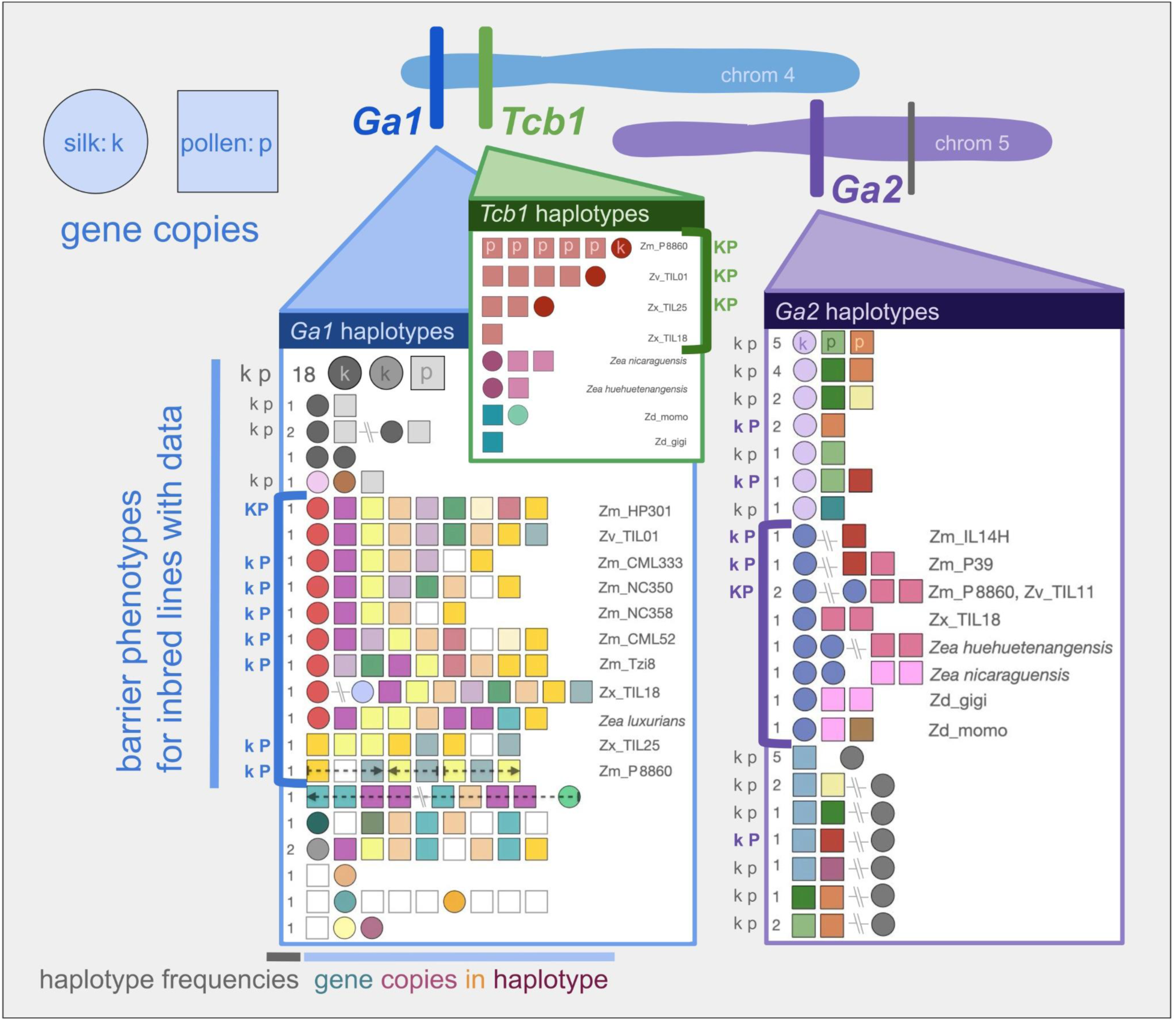
Haplotype diversity at each GA locus. Detailed view of haplotypes at the *Ga1* (blue) and *Tcb1* (green) loci on maize chromosome 4 and the *Ga2* (purple) locus on maize chromosome 5. Each row of shapes represents a sequence of full-length gene copies on a haplotype from 5’ to 3’, where circles are silk genes and squares are pollen genes. Shared gene copy color represents shared gene copy sequence. Gene copies with stop codons are gray, and gene copies found in only one haplotype are white. Hashed lines between gene copies represent more than a kilobase of distance between copies. Location of the distal non-functional silk gene copy found in some *Ga2* haplotypes is also marked on chromosome 5 in gray. Gene copies on a haplotype are all in the same direction, except where indicated by arrows. Haplotype frequency is labeled in gray to the left of each haplotype, except for *Tcb1* haplotypes which are all singletons. Known barrier activity is marked for inbred lines phenotyped for both silk (k) and pollen (p) activity, where bolded, colored, capital letters represent an active barrier (K) or ability to overcome the barrier (P). Haplotypes which seem similar to fully active haplotypes (indicated by brackets) are labeled with species and line names. Condensed species labels for genotypes with line names are: Zm (*Z. mays mays*), Zv (*Z. mays parviglumis*), Zx (*Z. mays mexicana*), and Zd (*Z. diploperennis*).

Out of the three GA loci, *Ga2* exhibits the least haplotypic variation across genomes, and the locus is present in all genomes we studied. Across *Zea* genomes, many different haplotypes are present at low frequency, as expected under a neutral model of allele frequencies (Ewens sampling distribution with θ=20; multinomial test *p*-value = 0.337831) [Supplemental table 9].

Unexpectedly, we discovered that a group of maize haplotypes missing a functional silk gene in the canonical *Ga2* locus is associated with an unmethylated, full-length copy of the *Ga2* silk gene nearly 50 Mb downstream of the *Ga2* locus [Supplemental table 3]. Each of these genomes shows a pollen gene at the syntenic position, but we find no evidence of structural rearrangements or genome duplications that can explain the distal silk gene location. Every *Ga2* silk gene copy with available methylation data, including distal *Ga2k* copies and *Ga2k* copies with premature stop codons, is unmethylated in diploid plant tissue (at CG, CHG, and CHH - ‘unmethylated’ corresponds to UMRs identified in Hufford et al. 2021). *Ga2k* copies are expressed in a variety of tissues; including silk- and pollen-containing tissues and roots; with the exception of the *Ga2k* distal copies, which are not expressed in any tissue [Supplemental figure 5, Supplemental table 3]. Pollen genes exhibit variation in methylation in diploid tissues; a few *Ga2p* copies with stop codons are unmethylated, and about half of the full-length copies of *Ga2p* display TE-like methylation in leaf tissue, which is characteristic of some highly expressed maize pollen genes (Zeng et al. 2023; Zeng et al. 2024). All *Ga2p* gene copies are expressed in pollen-containing tissue, and some *Ga2p* copies are also expressed in seeds [Supplemental figure 5, Supplemental table 3].

*Ga1* haplotype diversity is dominated by one haplotype which is found in most maize lines. This conserved haplotype is composed of two silk gene copies and one pollen gene copy, all of which have premature stop codons. Many other haplotypes are present, but at low frequencies, and the overall frequency distribution deviates strongly from simple neutral expectations (θ=38; multinomial p-value=1e-07) [Supplemental table 9]. Maize lines with documented Ga1 silk and pollen function (*Ga1-S* allele) or just pollen function (*Ga1-M* allele) all share a *Ga1k* gene copy but vary in *Ga1p* gene copy number and identity. Although silk expression data is not available for all *Ga1-S* lines, the silk gene in each of these lines is unmethylated and in a region of open chromatin. Pollen gene copies in these lines show TE-like methylation in leaf tissue, but are unmethylated and highly expressed in pollen (Zeng et al. 2024). Four maize lines with the *Ga1-M* allele (CML333, NC350, NC358, and Tzi8) show variable numbers of functional copies of the pollen gene and a highly methylated, unexpressed, full-length copy of the silk gene with promoter and coding sequences identical to expressed copies [Supplemental table 3, Supplemental table 1]. In another *Ga1-M* maize line that we examined (CML52), the full-length silk gene copy is unmethylated but lacks the ATAC signal typical of open chromatin that is found at active silk gene copies in other lines [Supplemental table 3, Supplemental table 1]. Observed expression of both silk and pollen functional *Ga1* genes is limited to reproductive tissues – anther and tassel tissues for pollen genes and silk for silk genes [Supplemental figure 5, Supplemental table 3].

The *Tcb1* locus displays presence/absence variation across the *Zea* genus, and is absent in the vast majority of sequenced cultivated maize lines. The diversity of present *Tcb1* haplotypes reflects the species phylogeny, with sets of similar haplotypes shared within species. *Tcb1* allele frequencies match neutral expectation (θ=8; multinomial *p*-value = 1) [Supplemental table 9]. Although published methylation data is unavailable for any of the published genomes containing *Tcb1*, we inferred CpG methylation across all three loci using HiFi reads from young leaf tissue that we also used for our P8860 genome assembly (Hall et al 2022). Methylation at the *Ga2* and *Ga1* functional genes in P8860 are as expected – *Ga2k* is unmethylated while *Ga2p* and *Ga1p* both have CpG methylation – while at the active Tcb1 locus, the *Tcb1k* gene is unmethylated, and all five *Tcb1p* copies are methylated, which we expect based on the TE-like methylation we observed at active *Ga1p* genes. In lines with Tcb1 present, *Tcb1p* is expressed in the tassel, while *Tcb1k* shows expression in both root and reproductive tissue, similar to *Ga1k* [Supplemental figure 5, Supplemental table 3].

### Evolution of the gametophytic factor genes and loci

To better understand the genetic relationship among gene copies and loci, we built separate phylogenies from full length coding sequences (CDSs) for silk and pollen gene models at all three loci [Supplemental figure 6]. Silk and pollen tree topologies match each other, as expected for two genes which evolved with a shared function (Fryxell 1996). The species tree topology is also reflected in the gene trees, where within each locus, closely related genes are found in closely related species [Supplemental figure 6]. The gene trees show that observed GA haplotypes consist of variable combinations of multiple distinct gene copies, identified by monophyletic subclades in the gene trees [Supplemental figure 6], and that many haplotypes with the same total gene copy number exhibit differences in the identity of the gene copies [Figure 3].

An important exception is that, in both the silk and pollen gene trees, the GA genes found in the most common *Ga1* locus haplotype appear to be a conserved set of three full-length genes - two non-coding silk genes and one non-coding pollen gene - each with CDSs containing distinct and conserved premature stop codons [Figure 3, Supplemental figure 6].

These non-functional genes are surprisingly diverged from validated functional *Ga1* genes and have an origin that is older than the split between functional *Ga1* and *Tcb1* genes. Specifically, based on sequence divergence, we estimate that these three older non-coding genes are roughly 610K and 750K years old (silk) and 1.5M years old (pollen), while the *Ga1* and *Tcb1* functional genes split ∼190K years ago (silk) and ∼1M years ago (pollen) [Supplemental table 2] (Clark et al. 2005).

We tested for episodic positive selection in both the silk and pollen gene trees by using a branch-site random effects model to check for elevated values of positive selection (ω) on all internal branches leading to divergence between GA gene types (Smith et al. 2015, Kosakovsky Pond et al. 2020) [Figure 4]. Both trees include an outgroup clade of ten other functional maize PME genes, which represent all B73 maize PME genes that have the PME EC 3.1.1.11 and have been described in multiple papers. In the silk gene tree, the branch subtending the *Tcb1* silk genes shows significant change in selection (*p*-value = 0.024). In the pollen gene tree, the branch subtending all GA pollen genes and the branch subtending almost all *Ga2* pollen genes show significant change in selection (*p*-value = 0.00006 for both), as does the branch subtending all *Tcb1* pollen genes (*p*-value = 0.011). The pollen gene tree branch with the next most significant change in selection (*p*-value = 0.056) is the branch subtending all *Ga1* and *Tcb1* pollen genes.

**Figure 4:**
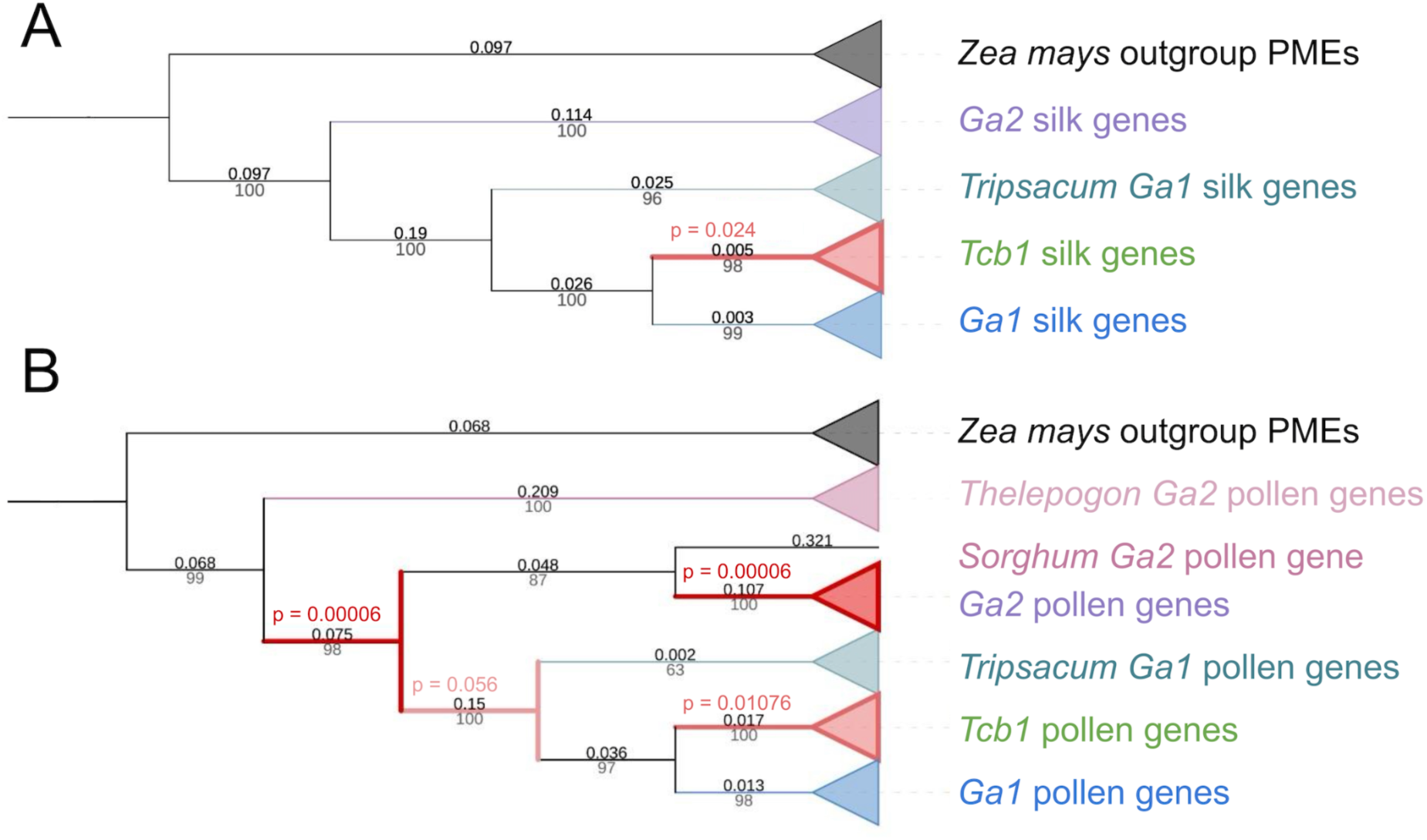
Positive selection during the evolution of functional GA genes. GA genes were under positive selection (ω) on gene tree branches subtending functional silk (A) and pollen (B) genes. Maximum likelihood trees were built in RAxML (Stamatakis 2014). Branches are labeled with lengths above in black and bootstrap values below in gray. Branches with significant evidence for positive selection are red, with additional p-value labels also in red on the (A) silk gene tree and (B) pollen gene tree.

### Molecular evolutionary analysis identifies patterns of constraint and adaptation on GA proteins

Because each GA locus generates a distinct barrier, we expect that positive selection resulted in specific amino acid changes which distinguish the barriers from each other. In particular, inactive pollen displays distinct morphology when growing down the silk depending on which silk-expressed gene – *Tcb1k*, *Ga1k*, or *Ga2k* – is active, so we expect the differences between GA silk amino acid sequences to drive the functional distinction between the barriers (Lu et al. 2014) [Figure 1]. To test for selection on individual amino acids that may control this impact of silk genotype on pollen growth, we used episodic positive selection tests on site changes from branches splitting the silk genes into distinct GA types (Murrell et al. 2012; Kosakovsky Pond et al. 2020). We identified 10 codons under significant positive selection on these branches of the silk gene tree (*p*-value < 0.1, the recommended *p*-value threshold for this test) [Supplemental table 4]. Because each GA silk protein interacts directly or indirectly with a paired GA pollen protein, we also checked whether the corresponding GA pollen genes may have evolved in concert. We found that 22 codons are under positive (*p*-value < 0.1) selection on the pollen gene tree [Supplemental table 4].

Notably, one of the four active site residues predicted to catalyze the PME reaction corresponds to a codon under positive selection on the branch subtending all pollen GA PMEs, where all outgroup and silk GA PMEs have a Q and all pollen GA PMEs have an E [Figure 5, Supplemental table 4]. This shift swaps the ancestral glutamine (Q) for a novel glutamic acid (E); this shift maintains the spatial volume but shifts the charge within the active site. While the activity shift caused is unclear, this is the only internal amino acid site under selection and is monophyletic for the change to GA pollen genes. Future work will need to ascertain its functional significance. Surprisingly, for both silk- and pollen-expressed genes, all of the sites under positive selection are on the surface of the predicted protein structures, where protein-protein interactions might occur. However, there is no overlap between the surface sites under positive selection and the sites where a known PME interaction with PMEI (Pectin methylesterase inhibitor) would occur (Di Matteo et al. 2005) [Figure 4].

**Figure 5:**
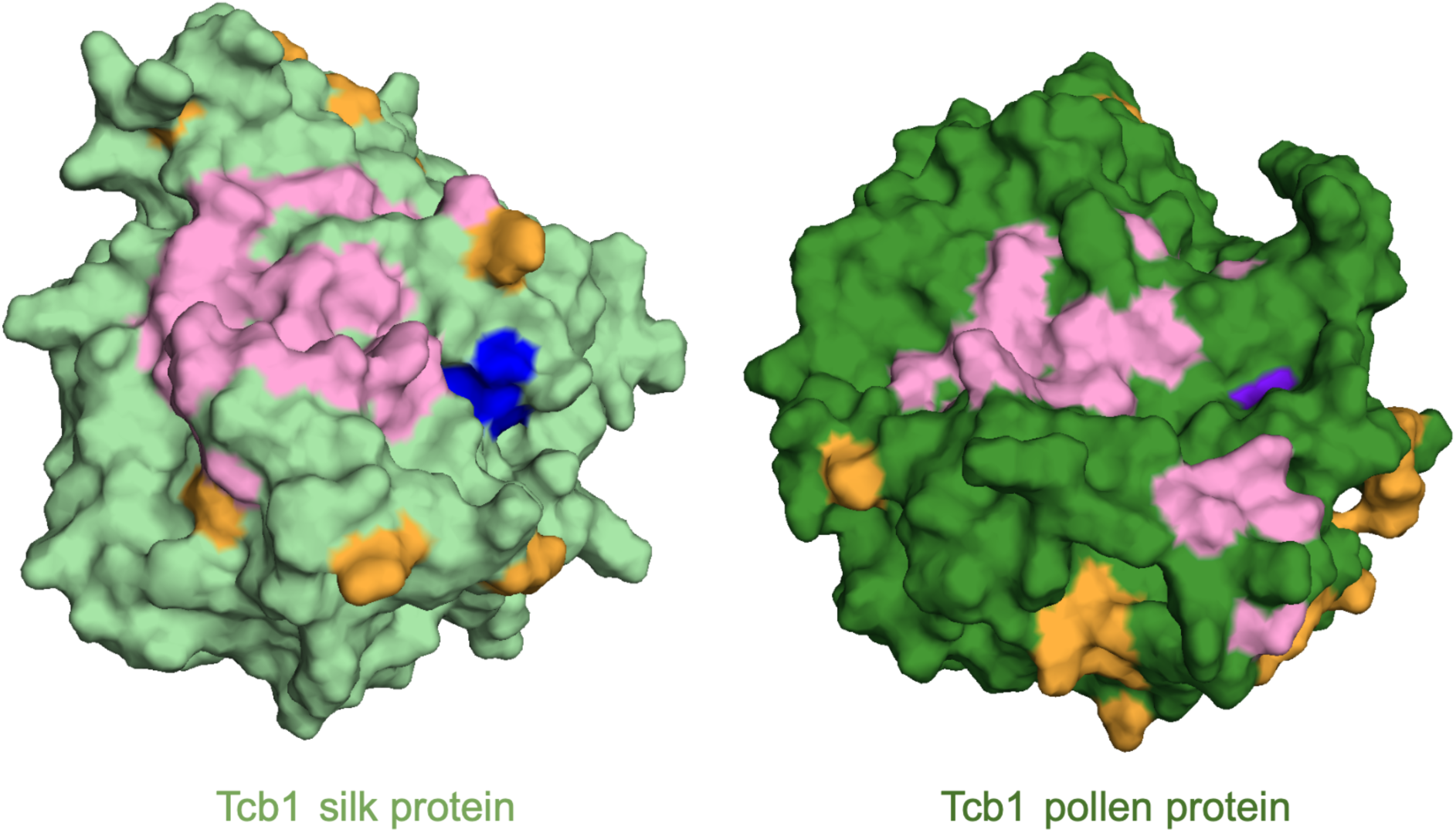
Sites under positive selection mapped onto 3D model of protein structure models of Tcb1 silk and pollen proteins. Colors represent active sites not under positive selection (blue), active site under positive selection (purple), other sites under positive selection (orange), and location of predicted PME-PMEI interaction surface (pink). Sites under positive selection are all on the surface and are not residues that participate in PME inhibition via PME-PMEI binding. Active site residues (blue) were identified via alignment to validated residues from (Johansson et al. 2002). PME-PMEI interaction residues are based on alignments to validated residues from (Di Matteo et al. 2005). Structures predicted with AlphaFold2 (Jumper et al. 2021) and visualized in PyMol.

### Lines with a conserved, inactive *ga1* haplotype are associated with specific 24-nt siRNAs in developing pollen

The conserved inactive *ga1* haplotype seems to serve some function; it is a haplotype that includes three highly conserved gene models that are in some cases expressed and unmethylated, and the haplotype is significantly more frequent than expected under a neutral model (see *Ga1* haplotype section). We propose that this haplotype is an *ga1* inactive allele with a non-barrier function, which we name *ga1-Off* (*ga1-O*). To investigate a potential non-protein-coding role of the *ga1-O* allele, we checked for an association between pollen siRNAs and the presence of this haplotype. Recent studies in maize have found pollen/anther-specific small RNAs may play an important role in pollen (Berube et al. 2024; Zhan et al. 2024). Using a database of siRNAs from the 0.4 mm (4 days prior to the start of meiosis in *Z. mays mays* when pollen is in an early mitotic stage of development) and 2 mm (late prophase I stage of meiosis in *Z. mays mays*) stages of anther development (Nakano et al. 2020), we searched for siRNAs in three maize inbred lines with *ga1-O* (B73, Oh43, and IL14H) and three genotypes with active versions of *Ga1* (*Ga1-S* maize inbred line HP301, *Ga1-M* maize inbred line NC358, and *Ga1-M mexicana* teosinte inbred line TIL25). We found that unique 24-nt siRNAs targeting reference GA silk gene sequences (*Tcb1k*, *Ga1k*, and *Ga2k*) are more abundant in 0.4 mm anthers of lines with the *ga1-O* allele compared to anthers with active *Ga1* alleles (*Ga1-S* and *Ga1-M*) (Welch’s t-test, *p*-value = 0.03633) [Figure 6]. In contrast, the number of siRNAs targeting reference pollen GA genes are similar across all genotypes (Welch’s t-test, *p*-value = 0.5238). Out of the *Ga1* lines included, only NC358, which is *Ga1-M* with a methylated *Ga1k* gene, had any of these silk gene-mapping siRNAs. Sequence comparison using BLAST reveals that many of the silk gene-mapping 24-nt siRNAs expressed in the *ga1-O* line B73 have SNPs unique to the *ga1-O* allele that are not found elsewhere in the B73 genome, indicating that the *ga1-O* allele is likely the source of 24-nt siRNAs and *Ga1* alleles the target [Supplemental figure 7] (Nakano et al. 2020). The 24-nt siRNAs we found to be enriched in the *ga1-O* lines are unphased [Supplemental figure 7]. In anther tissues, 24-nt siRNAs were the only length of siRNAs that showed a difference in number across genotypes, and the difference was only significant in the 0.4 mm and not the 2 mm (Welch’s t-test, p-value = 0.088) anther stage [Supplemental table 5]. In published siRNA data from maize inbred lines, 24-nt siRNAs mapping to the full-length silk gene copies in the *ga1-Off* region of the genome are present in 0.4, 0.7, 1.0, 1.25, 1.5, 2.0, 3.0, 4.0, and 5.0 mm long anthers, spanning thirty days of pollen development from the early mitotic to the binucleate microspore stage of pollen development (Zhai et al. 2015; Nakano et al. 2020). Phased 24-nt siRNAs generated by the somatic tapetal cells are transported into meiotic pollen cells (Zhou et al. 2022). Similarly, it may be possible that the unphased 24-nt siRNAs present in 0.4 mm anthers, during a stage of development before the tapetum has formed, may be generated by the somatic diploid anther tissues and either persist or continue being generated for weeks of anther development until at least the 5 mm stage, eventually impacting the developing haploid pollen (Zhan et al. 2024; Chow and Mosher 2023). We did not observe a difference in the number of siRNAs mapping to GA silk or pollen genes across genotypes in leaf or internode tissues (Nakano et al. 2020).

**Figure 6:**
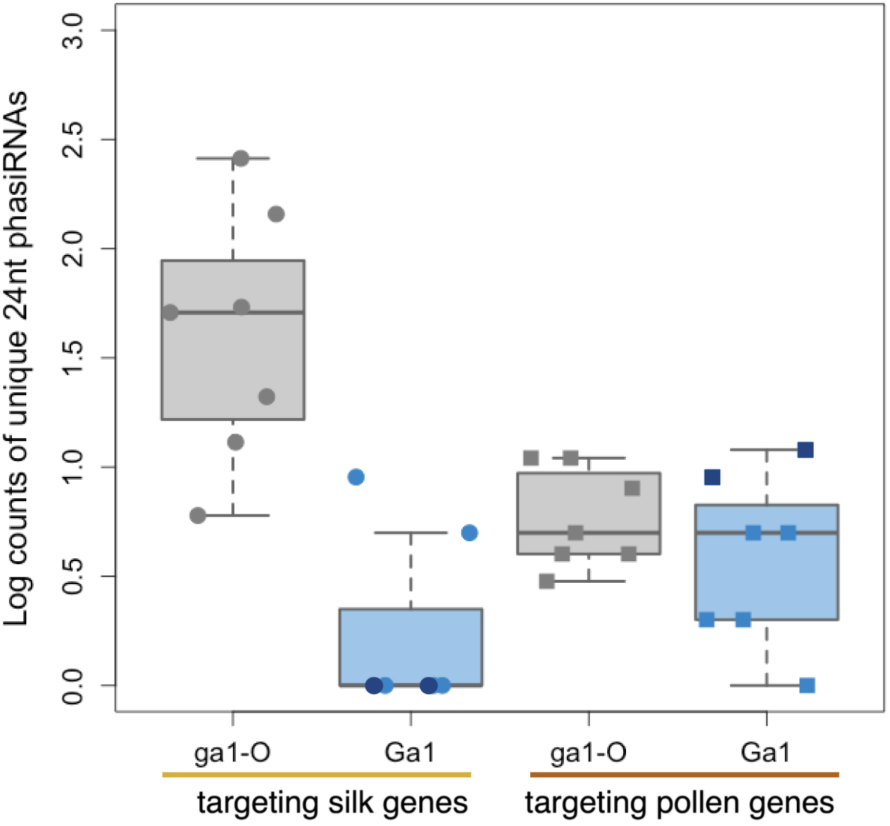
***Zea mays* pre-meiotic 0.4 mm anthers from *ga1-O* plants produce more unique 24-nt siRNA sequences targeting the GA silk gene sequences** than plants homozygous for *Ga1-S* (dark blue) or *Ga1-M* (light blue). There is no genotype-associated difference in the number of 24-nt siRNAs targeting GA pollen genes. Likewise, there are no genotype-associated differences in non-anther tissues.

## Discussion

### Gametophytic factors may have contributed to *Zea* diversification

The gametophytic factor loci *Ga2*, *Ga1*, and *Tcb1* generate reproductive barriers in *Zea mays* and have been proposed to be subspecies barriers (Evans and Kermicle 2001; Chen et al. 2022). Recent modeling work, however, suggests that the individual impact of any one of these loci is unlikely to prevent gene flow between species for more than ten thousand generations (Rushworth et al. 2022). This is partially due to the inability of GA-like loci to maintain reproductive isolation between distinct populations that come into contact, likely precluding their role in maintaining species boundaries (Rushworth et al. 2022). The presence of putatively functional copies of each barrier gene in more diverged genera, along with our sequence-based estimates of divergence times of individual genes, provides clear evidence that even the youngest of these loci, *Tcb1*, did not evolve recently to prevent gene flow from domesticated maize into teosinte.

One possible role for the Tcb1 barrier could have been to maintain distinction between diverging populations during the *Zea* genus diversification. Both the date of the *Tcb1k* origin around the time of *Zea* genus diversification, as also observed by Bapat et al. (Bapat et al. 2023; Chen et al. 2022), and the pattern of Tcb1 diversity across species are consistent with this role. *Tcb1k* is also completely absent in maize lines that do not generate a barrier; therefore, selection on this gene likely reflects selection on the functional barrier. Additionally, we observe positive selection on the branch leading to *Tcb1k* genes, suggesting that the role of Tcb1 as a species barrier was under selection before and as *Zea* was diversifying [see Figure 4]. Our observation of a full-length putatively functional copy of the *Tcb1k* gene in *Z. nicaraguensis* is evidence that the Tcb1 barrier may currently play a role outside of the *Z. mays* subspecies.

Conflicting with a possible role for Tcb1 barrier in species diversification via reproductive isolation, the age of *Tcb1p* is relatively ancient; we estimate *Tcb1p* diverged from *Ga1p* around 1M years ago, significantly predating the diversification of the genus ∼170K years ago (Chen et al. 2022b). Our relatively old age estimate for Tcb1p is based on synonymous substitution rate, which may be artificially elevated due to recent positive selection on non-synonymous and linked synonymous sites; for example, multiple sweeps rather than age could explain the high divergence of Tcb1p from Ga1p [Figure 4]. Further research is needed to functionally validate these and other putatively functional GA gene copies found in non-*Z. mays* genomes. However, the fact that functional silk-expressed gene copies are shared between *Z. mays* and other *Zea* species at all three barrier loci strongly supports the idea that these barriers are functional across the genus.

### Potential origins of the gametophytic factor reproductive barriers

The anatomy of maize and related grasses may have facilitated the retention of pollen-pistil barrier loci. Compared to related taxa, *Tripsacum* and especially *Zea* species have unusually long silks (stigmas) [Supplemental figure 8, Supplemental table 6]. After pollen germinates on a silk, the pollen tube grows into and down the remaining length of a transmitting tract inside the silk tissue to reach the female gamete. In species with longer transmitting tracts, this could present more opportunity for the pollen tube growth pattern and rate to become an important driver of fitness. Consistent with this idea, after the origin of the Ga2 barrier, duplicates like those which led to Ga1 and Tcb1 seem to only have been retained in the sister genera *Zea* and *Tripsacum* and not in related species with shorter silks. Long silks may have enabled GA barriers to evolve regardless of whether or not GA barriers conferred an adaptive benefit; the barriers are transiently reinforcing (Rushworth et al. 2022), so they may have been able to persist on short timescales just by selfishly excluding pollen from outside populations.

Long silks may have also increased the impact of pollen-silk interactions, including interactions between PMEs and other proteins, on fitness. One possible interaction protein could be a linked and silk-expressed pathogenesis-related protein called *ZmPRP3*, which has been proposed as a third component of the *Ga1* locus that enables pollen tube growth (Wang et al. 2022). This would suggest a potential overlap with the role of silk PMEs in mediating pathogen response, and introduces the possibility that the GA PMEs may have originally played a role in impeding pathogen growth down the silk (Begcy et al. 2024). However, to date no research has shown a role of GA loci in silk pathogen resistance, and *ga1* silks were not more susceptible to silk-invading fungal pathogens than *Ga1* silks (Begcy et al. 2024).

The GA barriers are often compared to pistil-pollen self-incompatibility mechanisms, and it has been suggested that the GA loci have origins in an ancestral self-incompatibility function (Kermicle and Evans 2005; Dresselhaus et al. 2011). Our results do not clearly support this hypothesis. Although many haplotypes have GA pollen genes and no corresponding GA silk gene, this is not evidence that the pollen function evolved first; we expect there to be strong selection against the opposite configuration of only a functional silk gene and no functional pollen gene, which would lead to incompatibility with all other plants. Better evidence for the ancestral function of these genes comes instead from expression patterns of the current functional gene copies. In both maize and *Sorghum bicolor*, we see expression of the *Ga2* silk-expressed gene in both the silk/stigma and in the root [Supplemental table 3] (see methods). Pectin methylesterases play an important role in cell wall formation, growth, and maintenance, and diverse PMEs are present in organisms as distantly related as bacteria and plants (Markovic and Janecek 2004; Shin et al. 2021). Although various PMEs work in concert to coordinate cell wall integrity in growing plant tissues, the age of the PME family means that these proteins are in many cases distantly related to each other despite their shared protein function. The silk and pollen PMEs encoded by the gametophytic factors are not closely related. In general, the exact mechanism of the interaction between GA PMEs is unclear, but there is no evidence supporting the idea that self-incompatibility was a role for these PMEs as they evolved.

Further complicating interpretation of the evolution of the GA silk and pollen PME interaction is the fact that interaction surfaces of the GA PMEs seem to have been under selection [Figure 4]. Although direct interaction of two PMEs has never been documented, PMEs often bind to PME inhibitors (PMEIs), which typically include a functional PME domain and an inhibitor domain (Di Matteo et al. 2005). To date, other direct interactions between PMEs and other types of plant proteins have not been documented. The prevalence of PMEs and PMEIs in the silk and pollen tube provides the opportunity for many PME-PMEI interactions, including in complexes of more than one PME and PMEI. For example, previous research has implicated an additional maize PME, *ZmPME10.1*, as a component of a complex in which the *Ga1* and *Ga2* silk and pollen PMEs interact (Chen et al. 2022; Zhang et al. 2018). Additionally, interactions of the GA PMEs with other proteins may be important. This is supported by the fact that all sites we found to be under positive selection are surface sites, but none overlap with the predicted site of PME-PMEI interaction.

### The *ga1-O* allele may function to suppress active gametophytic factors

We were surprised to find that the most common haplotype of the *Ga1* region, which we call the *ga1-Off* allele, is an inactive *ga1* haplotype present in lines which up until now have been considered to be fully inactive because they have no Ga1 barrier function [Figure 3].

Within this haplotype, the putative coding sequences of the three full-length gene models, including stop codons, are highly conserved despite the fact that all three models seem to be non-coding. All three genes diverged from *Ga1* between 0.6-1.5M years ago, well before the *Zea* genus diversification, and before the divergence between *Ga1* and *Tcb1*. The high frequency of the *ga1-O* haplotype in the population strongly suggests that this is not the result of a recent expansion of a neutral allele (p-value=1e-07, see Results). Instead, we suggest the genes are conserved because this haplotype likely functions as an allele which can suppress active silk gametophytic factors.

The identification of a *ga1-Off* allele helps to explain prior observations that an active *Ga1* barrier allele can be suppressed when crossed into certain backgrounds. In two previous studies, the action of a popcorn-derived *Ga1-S* allele conferred differing barrier strength after backcrossing into different inactive *ga1* backgrounds (Nelson 1952; Ashman 1975). *Ga1-S* shows dominance when introduced into maize dent inbred Hy, but the barrier strength is significantly reduced when introduced to two different popcorn inbreds, Sg1533 and Sg18 (Nelson 1952; Ashman 1975). Using SNP data from the Ames 282 panel, we found that Sg1533 and Sg18 likely both carry the B73-like *ga1-O* allele, while Hy carries a B97-like *ga1* allele (Flint-Garcia et al. 2005; Bukowski et al. 2018) [Supplemental figure 9]. The B97-like *ga1* allele, in contrast to the *ga1-O* allele, is not notably common nor conserved and seems to behave as a truly non-functional ga1 allele, so we call this *ga1-Null* or *ga1-N*. We argue that the reduced activity of the *Ga1-S* allele in these experiments can be ascribed to the *ga1-O* v *ga1-N* identity of the inactive *ga1* allele in a heterozygous background. *ga1-O* gene copies are equally related to functional *Ga1* and *Tcb1* genes. [Supplemental figure 3, Supplemental table 2]. Consistent with this phylogenetic relationship, *ga1-O* seems to also silence active *Tcb1*. After 10 generations of backcrossing a *Tcb1-S* allele into W22, an inbred that carries the *ga1-O* allele, the teosinte Tcb1 barrier activity was fully suppressed in two independent lineages (Lu et al. 2014). When these lines with suppressed Tcb1 barriers were crossed into backgrounds lacking the functional *Mop1* RNA-dependent RNA polymerase, some of the offspring regained Tcb1 barrier function and *Tcb1k* expression (Lu et al. 2019).

Additionally, in sympatric populations of maize and teosinte, the Tcb1 barrier has only been observed in populations where *Ga1* is at least partially active (*Ga1-S* or *Ga1-M*) (Kermicle 2006; Kermicle et al. 2006) [Supplemental table 7]. Because *ga1-O* is by far the most common *ga1* allele and Tcb1 is active in most of these teosinte populations, the absence of Tcb1 barriers here may indicate that *ga1-O* is suppressing Tcb1 activity. It is possible that *ga1-O* may similarly regulate Ga2, based on sequence similarity to Tcb1 and Ga1, and 24-nt siRNA sequence match to *Ga2k* [Supplemental table 5], and anecdotal evidence of Ga2 barrier suppression. However, Ga2 and Ga1 barriers are active in populations with *ga1* alleles (Kermicle et al. 2006; Kermicle and Evans 2010) [Supplemental table 7], and we found no evidence of *Ga2k* genes being methylated in any background, so the Ga2 locus may be under a different type of regulation.

Future experiments will be required to establish a causal connection between the *ga1-O* locus and silk-expressed barrier function at any of the three gametophytic factor loci.

The exact functional and mechanistic difference between the two types of inactive *ga1* alleles we observed, *ga1-O* and *ga1-N*, is unclear from our experiments. However, Ashman suggested a similarity between what he termed the “suppression” of the *Ga1-S* allele and known maize paramutations (Ashman 1975). We also observe parallels between our findings and more current understandings of paramutation.

### The *ga1-O* allele and paramutation

In 1956, R. A. Brink coined the term “paramutation” to describe the heritable, but non-genetic, alteration of the *R* locus allele *R-r* when in a heterozygous plant with an *R-st* allele (Brink 1956). The altered allele, now *R-r’*, shows reduced expression of the red color associated with *R-r*. This heritable, but not permanent, silencing is enacted through methylation directed by the *R-st* allele (Walker 1998). Some shared attributes of well-studied paramutations in maize, as reviewed by Hollick, are as follows (from Hollick 2017): association with duplicated regions, locus specificity, transfer across generations, and implication of 24-nt siRNAs. The *Ga1* locus seems to largely share these attributes, though transfer of the paramutation across generations has never been tested. However, recent research proposes a model under which 24-nt siRNAs play a role in the RNA-directed DNA Methylation (RdDM) and control methylation associated with maize paramutations, which is consistent with all of our results (Deans et al. 2024).

The epigenetically silenced *Ga1k* genes may be thought of as carrying a paramutation. The methylated status of *Ga1k* genes in *Ga1-M* maize lines, and the possibility that *ga1-O* may be involved in a 24-nt siRNA-mediated RdDM pathway that could methylate GA silk genes both support this idea. Expanding the paramutation framework to the full locus, *ga1-O* acts as a paramutagen and *Ga1-S* (Ga1 locus with active silk and pollen factors) is paramutable. When *Ga1-S* is exposed to *ga1-O* in a heterozygous plant, *Ga1-S* may be epigenetically modified to become the paramutant *Ga1-S*’ allele (aka *Ga1-M* or a Ga1 locus with a silenced silk and an active pollen gene). This framework is consistent with our observation that demethylation and open chromatin are associated with active silk genes, while all *Ga1-M* maize haplotypes have silk genes that lack demethylation, open chromatin, or both [Supplemental table 3]. Similarly, some of the *Ga1-W* (weak barrier), *Tcb1-W* (weak barrier), and *Tcb1-M* alleles may also be thought of as *Tcb1-S’* alleles [Table 1]. While Ga1 has never been classified as a paramutation during the past century of GA research, this may reflect the fact that the barrier phenotype is much harder to observe than the visual color phenotypes associated with all but one known maize paramutation (Hollick 2017; Pilu et al. 2009). Without linked marker genes, even active barriers may be hidden from observation in an open-pollinated field.

Importantly, we have not established a strong causal link between the *ga1-O* locus and methylation of silk *Ga1* genes, nor have we established a causal link between methylation of silk Ga1 genes and activity of the barrier. To determine whether Ga1 is truly a paramutation system would require rigorous testing of the behavior of *Ga1-S* and *ga1-O* alleles in different backgrounds and across generations.

## Methods

### Genome assembly of maize line P8860

Maize line P8860, which creates and overcomes the Tcb1 and Ga2 barriers and overcomes the Ga1 barrier, was provided by Jerry Kermicle. High molecular weight DNA was extracted from young leaf tissue and sequenced via HiFi long-read sequencing on a Pacific Biosciences Sequel II at the UC Davis Genome Center. Reads were then assembled into a reference-quality genome using the Hifiasm assembler (Cheng et al. 2021), manually curated, and scaffolded using ALLMAPS (Tang et al. 2015). The P8860 genome assembly consists of 1,105 contigs with a mean contig length of 2,080,026 bp and a contig N50 of 91,160,284 bp for an estimated 99.90% coverage of the 2,300 Mbp genome. Genome features were annotated based on homology, resulting in complete versions of a total of 98.22% of the Poales obd10 BUSCO and 93.97% of the Liliopsida obd10 BUSCO genes (Manni et al. 2021; Manni et al. 2021). CpG methylation was then inferred using the same HiFi reads and the software primrose (Hall, R., Portik, D., Nyquist, K., & …), developed by Pacific Biosciences (accessed October 2022), which has since been replaced by an updated version of this tool called Jasmine (https://github.com/pacificbiosciences/jasmine/). Genome assembly and annotation will be made available at time of publication.

### Gene identification

We used previously published and validated versions of the *Tcb1k*, *Ga1k*, *Ga1p*, and *Ga2p* genes as our reference version of each GA gene type, but the published *Ga2k* and *Tcb1p* gene references did not appear in the genome assemblies of maize lines we knew had full *Ga2* and *Tcb1* activity (Lu et al. 2019; Zhang et al. 2023; Moran Lauter et al. 2017; Zhang et al. 2018; Chen et al. 2022). To identify improved versions of the *Ga2k* and *Tcb1p* gene models, we searched within the *Ga2* and *Tcb1* regions of the P8860 genome for full-length gene models similar to the known GA silk and pollen genes, respectively. We identified new *Ga2k* and *Tcb1p* reference genes (see Results and Supplemental figure 1 for comparison between published gene models and our two reference gene models). These reference genes are present in all genome assemblies from plants known to create the Ga2 barrier and overcome the Tcb1 barrier, respectively. The new reference genes include signal peptides, have the conserved one-intron structure for GA PMEs, and are expressed in plants and tissues known to create and overcome the barrier [Supplemental table 3].

To identify GA genes across diverse Andropogoneae genomes, we BLASTed for GA reference gene sequences against the NAM v5 maize genomes, 4 Teosinte Inbred Lines (TILs), the 30 PanAnd project genomes, maize line Mo17, maize line W22, and our new genome assembly of maize line P8860 (Hufford et al. 2021; Woodhouse et al. 2021; Edward Buckler Elizabeth Kellogg Adam…). We searched specifically for gene models supported by transcriptome annotations where available. We recorded genomic coordinates and pulled out CDS nucleotide and amino acid sequences for all high-quality hits (almost 100% match to the reference sequence used in our search query). In general, our BLAST hits for *Tcb1* and *Ga1* genes almost completely overlap due to similarity between the two loci, so we sorted *Tcb1* and *Ga1* hits by genomic location. We determined the genomic location of each locus by restricting just to the region between the area syntenic to the maize genes on either side of the Tcb1 and Ga1 loci on the maize consensus genetic map (available on MaizeGDB, accessed in 2023).

### Gene alignment and tree building

We aligned GA silk and pollen genes separately due to sequence dissimilarity between the two types of PMEs. To compare GA PMEs to Zea mays PMEs more broadly, we also included maize reference PMEs as an outgroup in all alignments and trees. We chose reference PMEs by searching the B73 genome annotation for genes with the Enzyme Code 3.1.1.11 for PME activity, then selected a subset of those genes which were annotated on MaizeGDB as being cited multiple times in published literature. We aligned these validated reference genes to each other, and we removed two PME genes that were too structurally different to be considered a useful reference. This resulted in 10 well documented reference PMEs that were used for comparative alignments [see Supplemental figure 1 for a list of these genes].

To focus the alignment on just the part of the gene that can be subjected to tests for positive or negative selection, we restricted the alignment to the CDS. Additionally, because signal peptides are under different selective pressures and evolve at a different rate than the amino acids that make up the mature protein, we used TargetP 2.0 to predict and cleave signal peptides at the beginning of all GA PMEs (Almagro Armenteros et al. 2019). We then aligned cleaved CDSs for all IDed gene models using Muscle5 with default parameter settings (Edgar 2022). Sequence alignments with all gene models, including those with stop codons in their CDS, were aligned without respect to codon position. Sequence alignments for only functional gene models, identified as those with identical or very similar sequence to gene models found in genomes from lines with GA silk or pollen function in the barrier phenotype, were also aligned without respect to codon position. Using cleaved CDS alignments, we assembled all gene trees in RAxML under a gammagtr model and bootstrapped all trees with at least 100 trials (Stamatakis 2014).

### Expression and methylation data

For all identified *Zea* GA gene models on main chromosomes (not scaffolds or contigs), we checked expression, methylation, and open chromatin (ATAC-peak) status (Hufford et al. 2021; Woodhouse et al. 2021; Edward Buckler Elizabeth Kellogg Adam…) [Supplemental table 3]. We also checked the expression of the *Sorghum bicolor* Ga2 gene models, as this is the one other genome in our study with a wide range of publicly available RNAseq data aligned to a reference genome (Moreno et al. 2022). The ortholog of *Ga2k* (SORBI_3004G350500) is expressed in the inflorescence, seed, and drought-stressed root, while the ortholog of *Ga2p* (SORBI_3004g350400) is expressed in the inflorescence, anther, and pollen (Varoquaux et al. 2019; Davidson et al. 2012; Wang et al. 2018).

### GA loci age estimation

To estimate the age of the GA loci, we separately estimated the divergence times for functional silk and pollen copies of *Tcb1*, *Ga1*, and *Ga2*. Using MEGA, we ran a K2P model to calculate synonymous substitution rate between pairs of cleaved CDSs (Tamura et al. 2021; Stecher et al. 2020; Nei and Kumar 2000) [see Supplemental table 2]. We then used this substitution rate to calculate the divergence time by dividing by the two branches coming off of the shared ancestral node between the gene pairs and a constant average maize mutation rate (Clark et al. 2005), giving an estimate of generation time since divergence. Since maize is an annual species, we assumed that generation time was 1 year, and converted generation time to years. We compared pairwise generation times of functional genes from each locus, and averaged across unique pairwise comparisons to get an average divergence time between each type of gene that came from a duplication of a previous version of the locus (eg. *Tcb1k* and *Ga1k* v *Ga2k* for the *Ga1* silk gene age estimate, and *Tcb1k* v *Ga1k* for the *Tcb1* silk gene estimate). For all pairwise comparisons, see Supplemental table 2.

### Neutral allele frequency test

To test for evidence of selection at each locus, we compared the observed haplotype frequency spectrum to that expected under a simple neutral model. Each present observed haplotype [Figure 3] was considered an allele in an observed allele frequency distribution [Supplemental table 8]. Expected allele frequency distributions were calculated for each locus using Ewen’s sampling distribution, with N representing the total number of genomes with present observed haplotypes of the locus, and the population mutation rate θ chosen via a grid search [Supplemental table 8]. We used a multinomial test implemented in R with the package EMT (Menzel 2010) to calculate a p-value for the comparison of the observed haplotype distribution to that expected under the maximum likelihood value of θ.

## Conclusion

For all three maize gametophytic factors, we documented haplotype and gene diversity, identified sites under positive selection, and estimated the timing of gene and locus divergence. We also sequenced a maize line with all three barriers at least partially active, which allowed us to observe a correlation between gene methylation and barrier activity at all three silk genes.

This silk gene methylation may be regulated by pollen-expressed 24-nt siRNAs created by the *ga1-O* allele. Future work would be needed to functionally validate the role of the *ga1-O* allele by establishing a causal relationship between the allele, the associated 24-nt siRNAs, and the silencing of the Tcb1, Ga1, and Ga2 barriers.

## Acknowledgements

We would like to thank J L. Kermicle for providing the P8860 line seeds and for invaluable feedback and advice on the direction of this project, C A. Rushworth for discussions about genetic conflict and evolutionary dynamics, and V Walbot for revelatory comments on a previous version of this manuscript. This work was supported by funding from the National Science Foundation (Plant Genome Research Project grant 1753632, Plant Genome Research and Plant-Biotic Interactions grant 2020754, and Systems and Synthetic Biology grant 1906486), the US Department of Agriculture (Hatch project CA-D-PLS-2066-H 548), and the UC Davis Plant Sciences Department (Graduate Student Research Award). This research used the High-Performance Computing Core Facility (HPCCF) at the University of California, Davis.

