## Supplemental Figures for "Molecular evolution of a reproductive barrier in maize and related species"

Supplemental figure 1: Alignments of published and new GA reference genes

Supplemental figure 2: Gene trees of putatively functional GA genes

Supplemental figure 3: Simplified gene trees with estimated divergence times

Supplemental figure 4: *Ga2* and *Ga1* are not duplicates from the Tripsacinae WGD

Supplemental figure 5: Expression anatograms by gene

Supplemental figure 6: Gene trees of all full-length GA genes

Supplemental figure 7: 24nt siRNA hits and phase scoring for each ga1-O ‘gene’ in B73

Supplemental figure 8: Stigma length and GA locus PAV on Andropogoneae species tree

Supplemental figure 9: SNP trees for GA loci across diverse maize lines

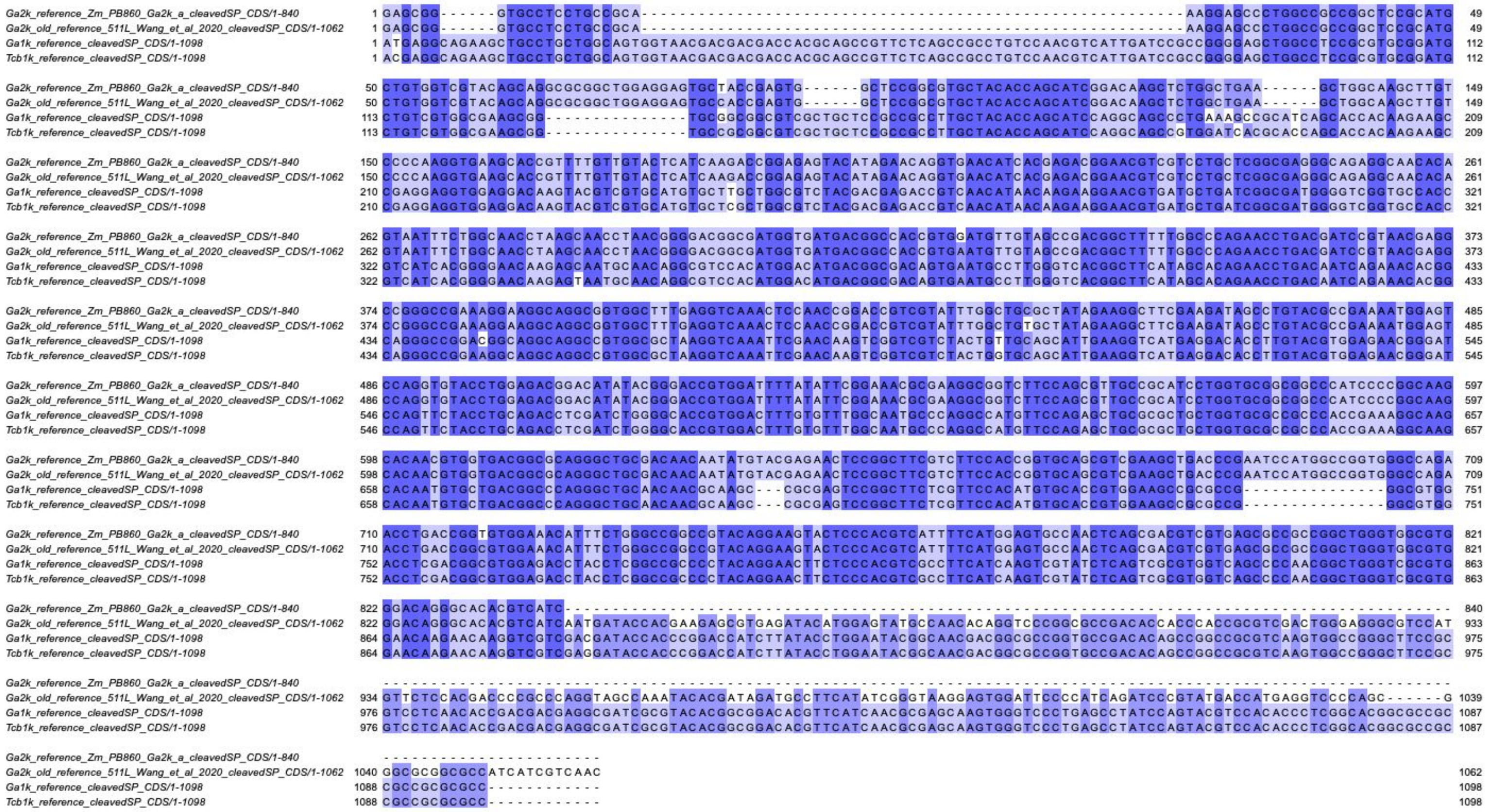


**Supplemental Figure 1a**
**Alignment of published and new GA silk reference gene CDSs**
Alignment of GA silk reference gene sequences. Our new *Ga2k* sequence has a shorter CDS. Codon-aware nucleotide alignments of nucleotide CDSs were made in muscle and visualized in Jalview. Color shows percent shared identity across sequences.

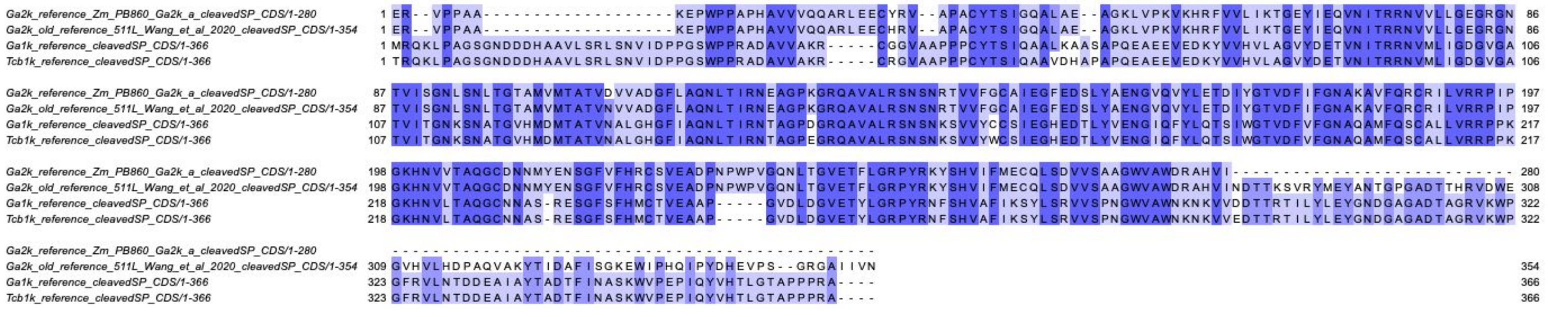


**Supplemental Figure 1b**
**Alignment of published and new GA silk reference gene protein sequences**
Alignment of GA silk reference gene amino acid sequences. Our new *Ga2k* sequence has a shorter CDS. Alignments were made in muscle and visualized in Jalview. Color shows percent shared identity across sequences.

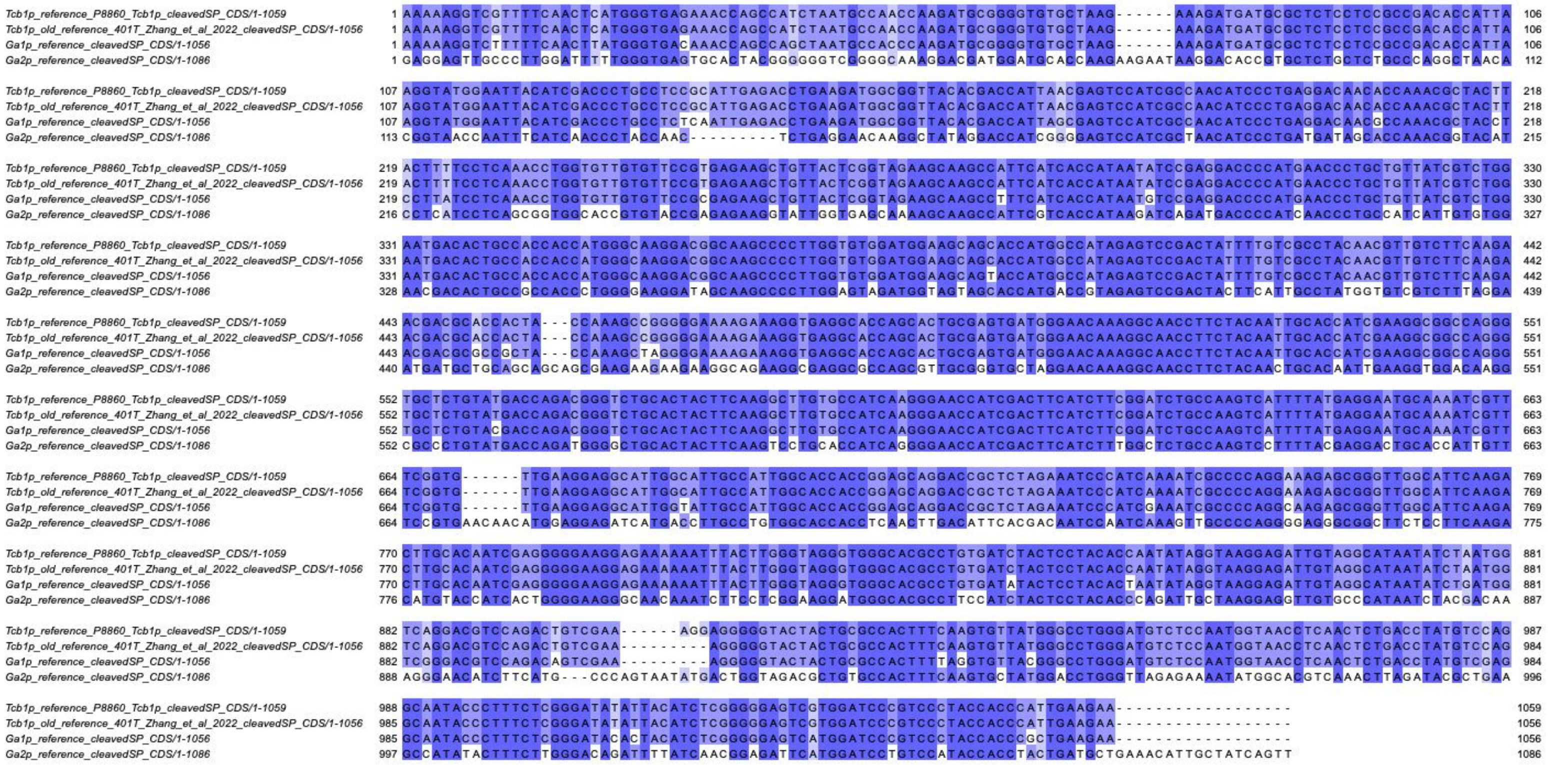

#### Supplemental Figure 1c
**Alignment of published and new GA pollen reference gene CDSs**

Alignment of GA silk reference gene sequences. Our new *Tcb1k* sequence has a shorter intron, which leads to a single codon change at nucleotide 904-906. Codon-aware nucleotide alignments of nucleotide CDSs were made in muscle and visualized in Jalview. Color shows percent shared identity across sequences.

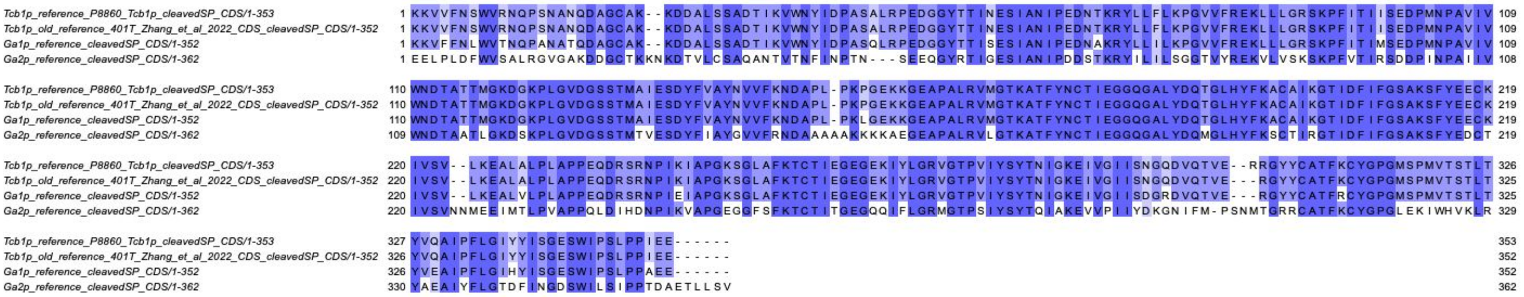

#### Supplemental Figure 1d
**Alignment of published and new GA pollen reference gene protein sequences**

Alignment of GA pollen reference gene amino acid sequences. Our new *Ga2k* sequence has an additional arginine at amino acid 302. Alignments were made in muscle and visualized in Jalview. Color shows percent shared identity across sequences.

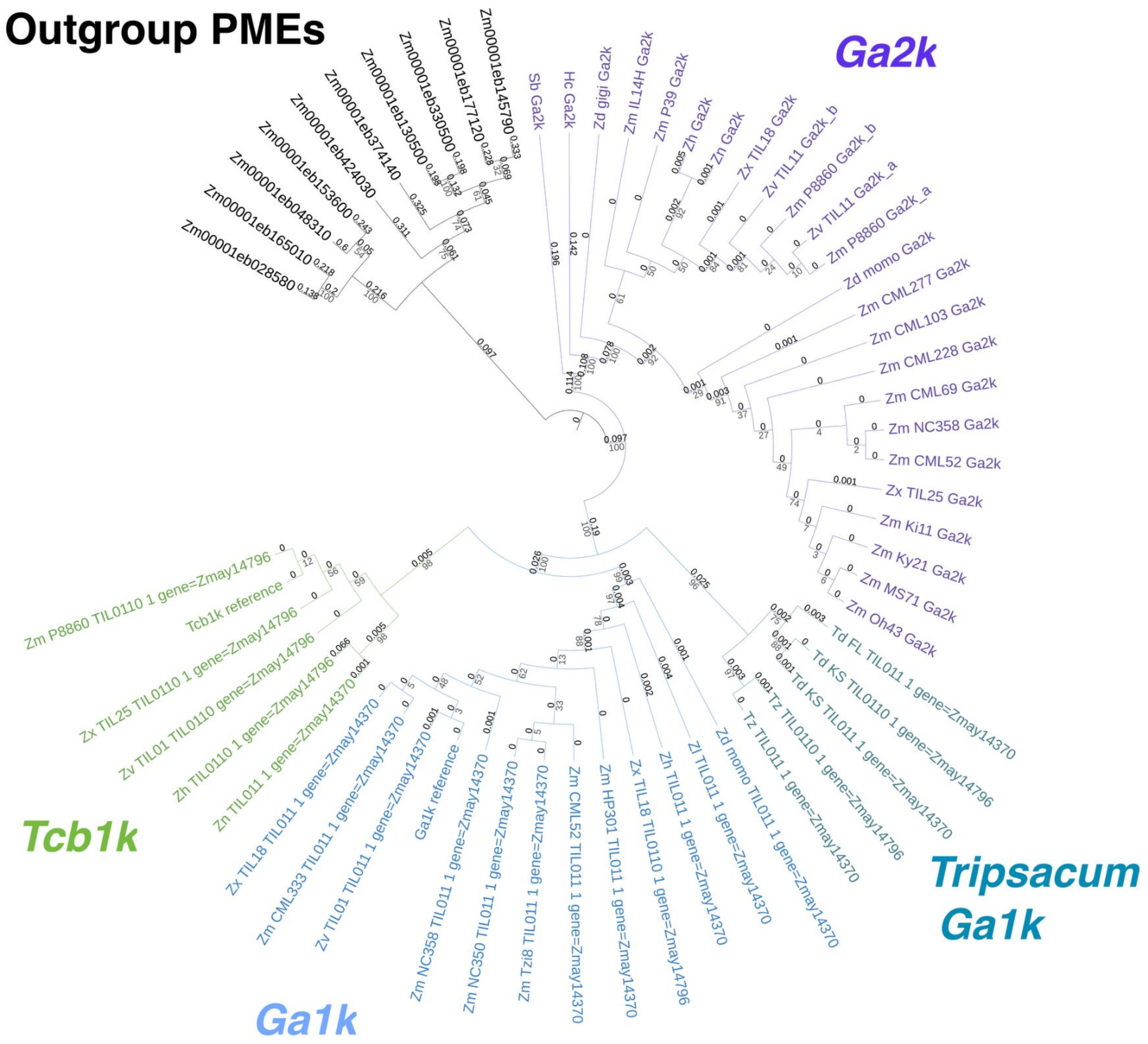


**Supplemental Figure 2a**
**Putatively functional GA silk gene tree**
Gene tree of GA silk gene sequences with CDS that can be translated into a full-length amino acid sequence with no premature stop codons. *Zea mays mays* PMEs as an outgroup. Tree is based on codon-aware nucleotide alignments of CDSs. Gene tree was built in RAxML and visualized in iTOL.

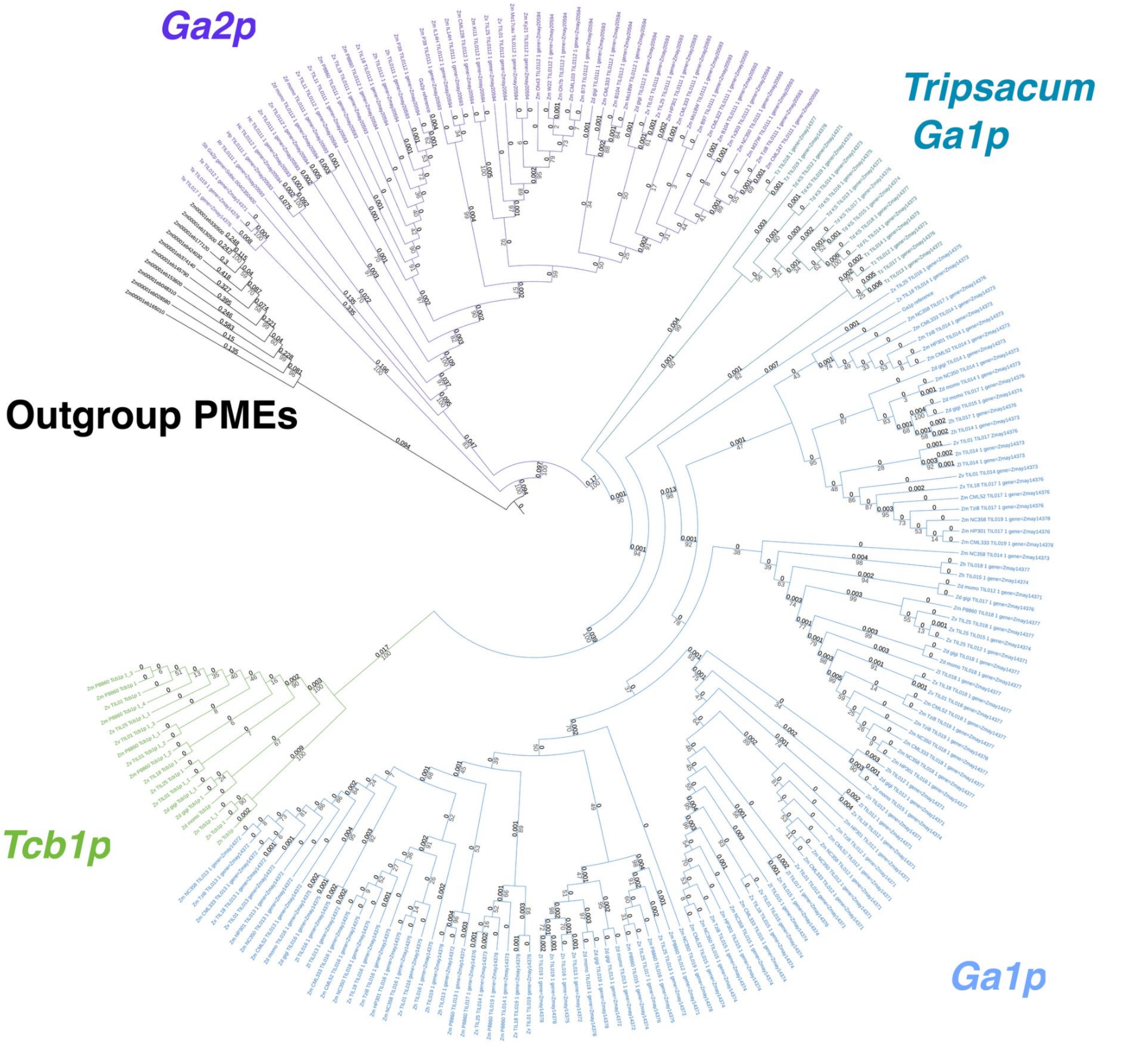


**Supplemental Figure 2b**
**Putatively functional GA pollen gene tree**
Gene tree of GA pollen gene sequences with CDS that can be translated into a full-length amino acid sequence with no premature stop codons. *Zea mays mays* PMEs as an outgroup. Tree is based on codon-aware nucleotide alignments of CDSs. Gene tree was built in RAxML and visualized in iTOL.

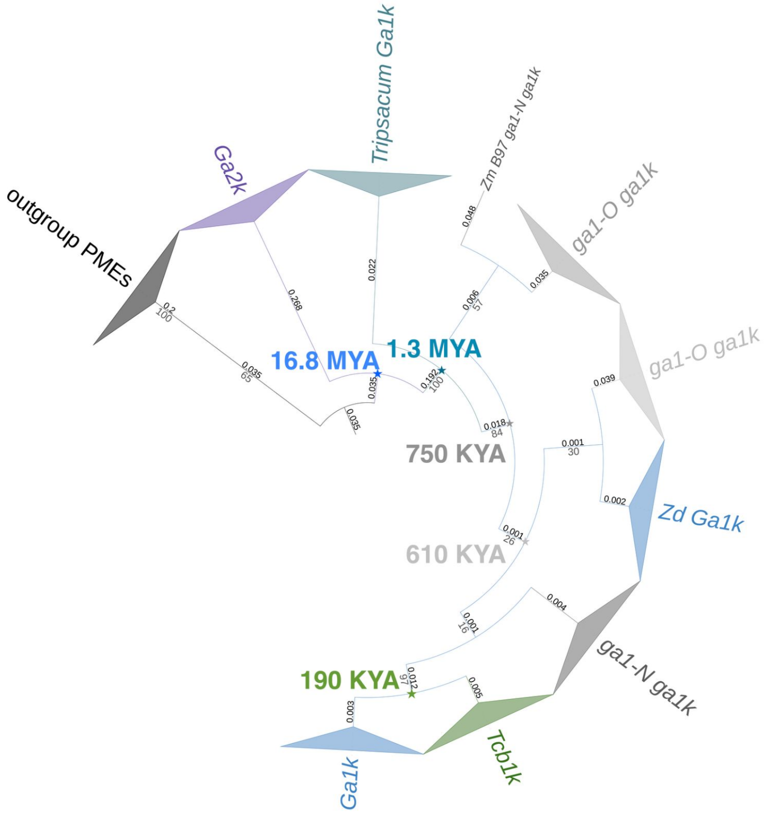


**Supplemental Figure 3a:**
**GA silk estimated divergence times**

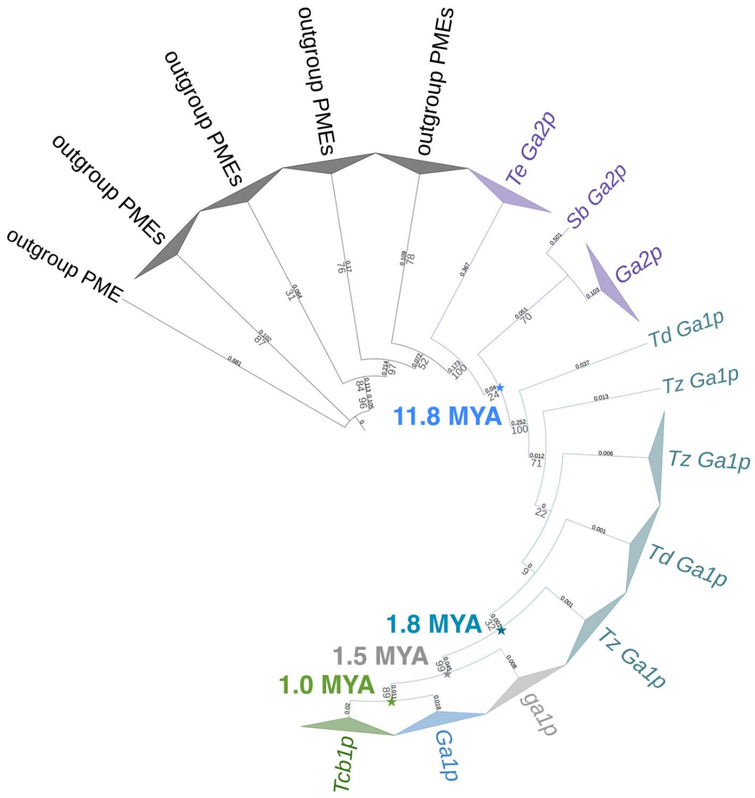


**Supplemental Figure 3b:**
**GA pollen estimated divergence times**

#### Supplemental Figure 3
#### Simplified GA gene trees with estimated divergence times

Gene trees simplified by locus with estimated ages. Ages were estimated via synonymous substitution rate and an assumed generation time of 1 year. *Zea mays mays* PMEs as an outgroup. Trees are based on codon-aware nucleotide alignments of CDSs. Gene trees were built in RAxML and visualized in iTOL.

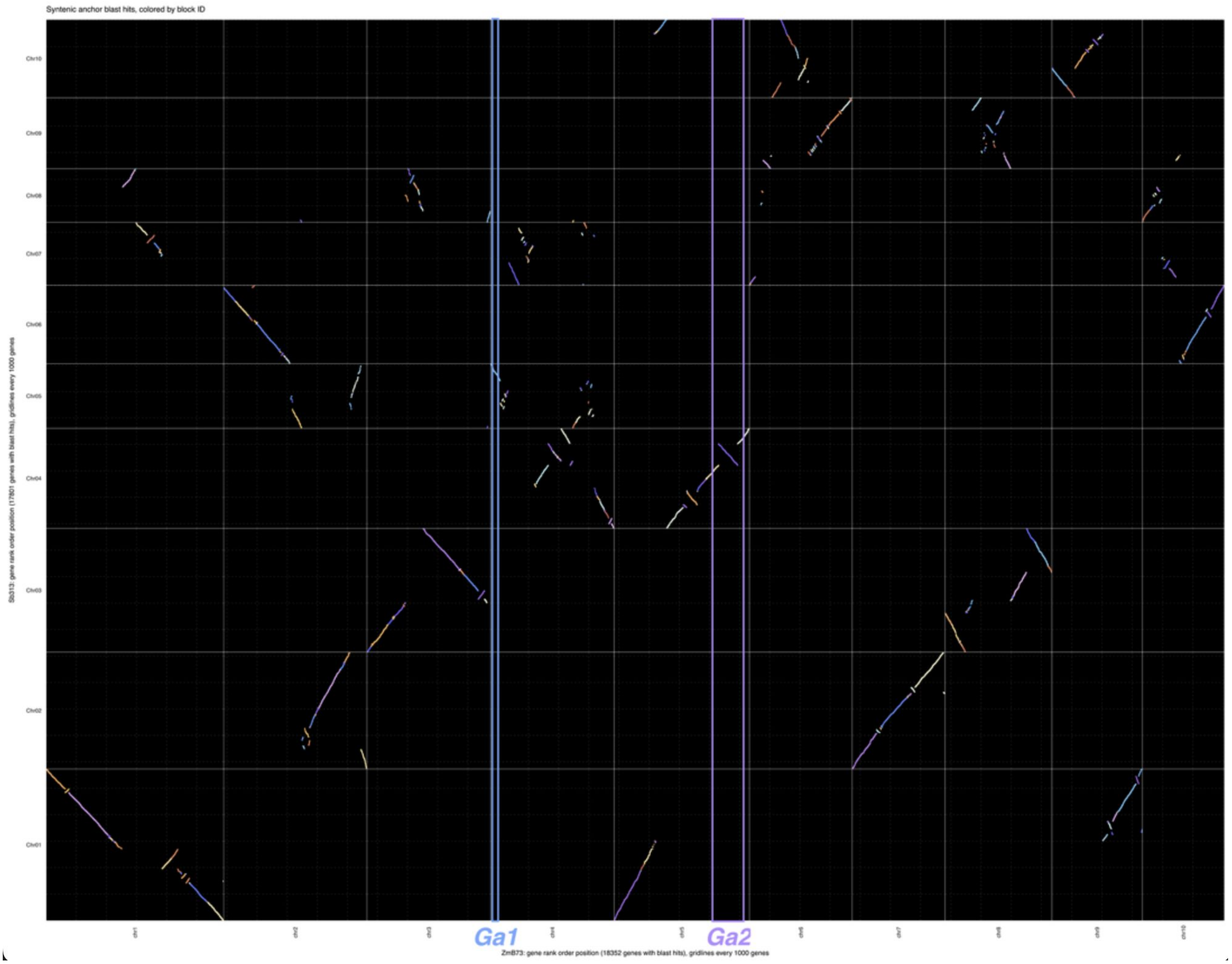


**Supplemental Figure 4**

***Ga2* and *Ga1* are not duplicates from the Tripsacinae WGD**

Whole genome synteny plot of orthologous genes between maize line B73 reference genome v5 and *Sorghum bicolor* reference genome v3.1.3. Sorghum and maize diverged before the Tripsacinae whole genome duplication, and fragments of the maize genome that arose during the WGD duplication are generally syntenic to the same region of the sorghum genome. Regions of the maize genome containing *Ga1* (blue box) and *Ga2* (purple box) are not syntenic to the same region of the Sorghum genome. Comparison generated in GENESPACE (Lovell et al 2022): https://doi.org/10.7554/eLife.78526

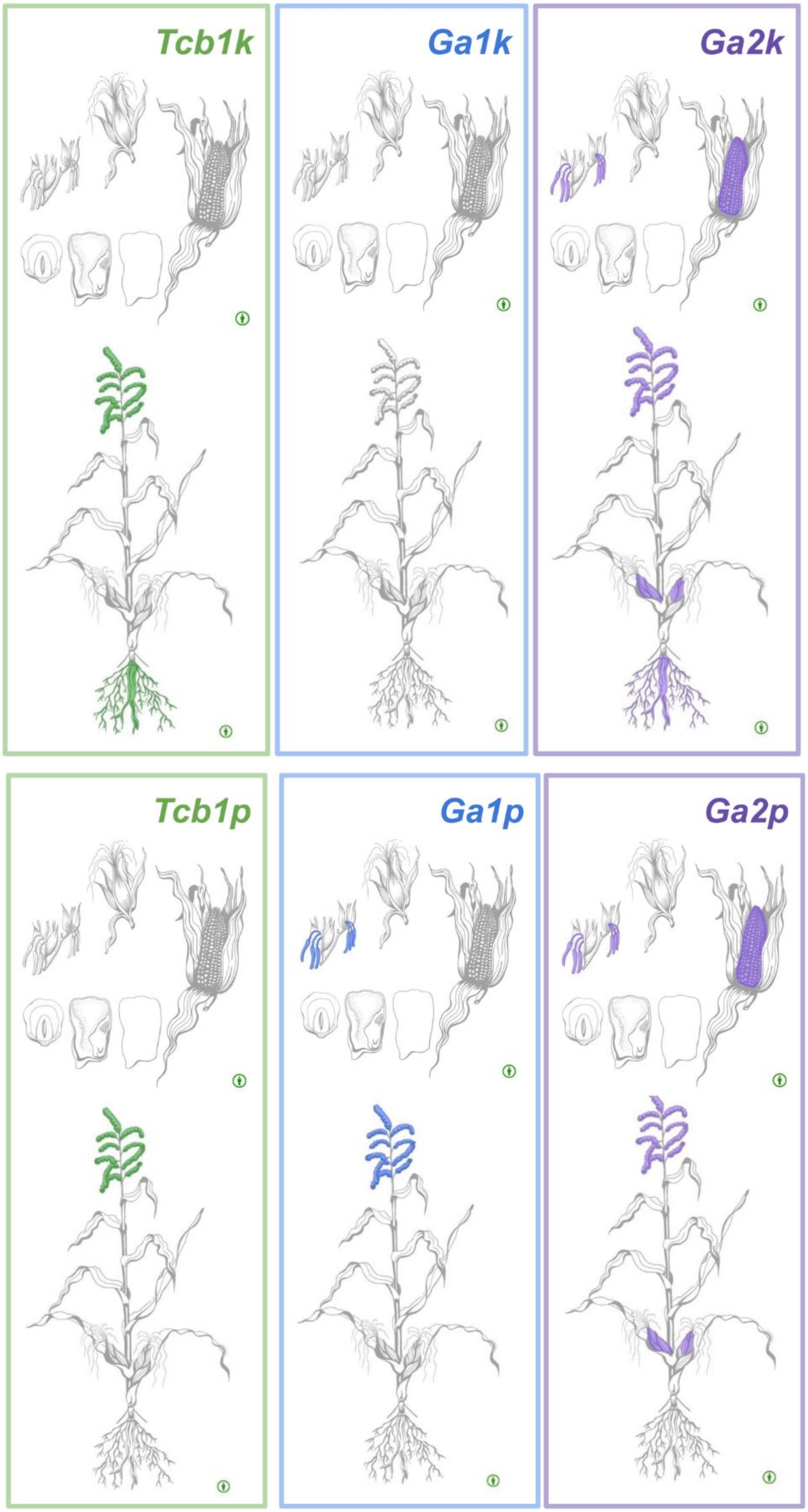


**Supplemental Figure 5**
**Expression anatograms by gene**
Tissues with RNA expression in at least one genome at >= 5 rpkm. See supplemental data for expression data sources. Anatogram templates are from Moreno et al 2022 https://doi.org/10.1093/nar/gkab1030.

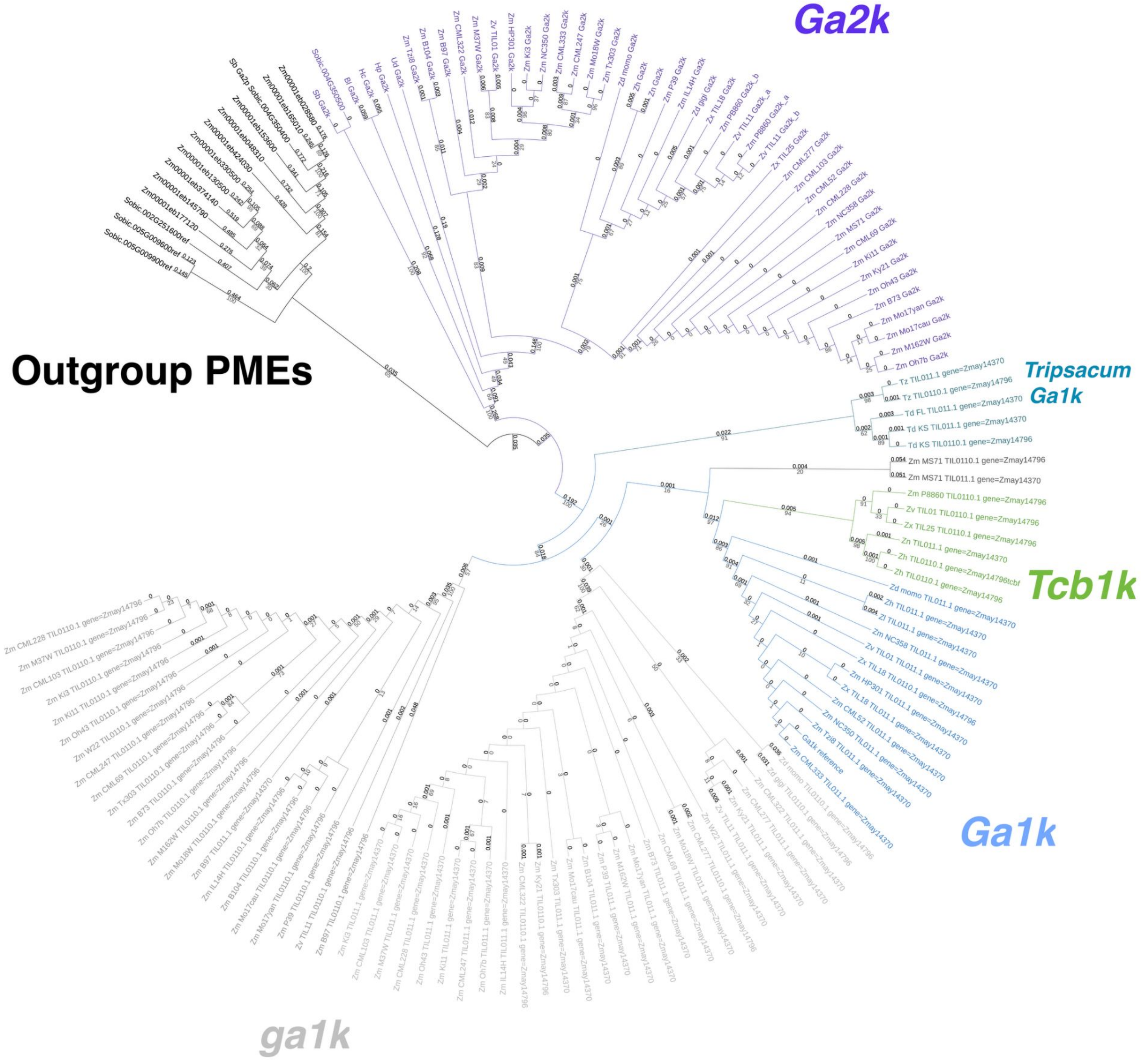


**Supplemental Figure 6a**
**GA silk gene tree**
Gene tree of all full-length GA silk gene sequences with *Zea mays mays* PMEs as an outgroup. Tree is based on nucleotide alignments of CDSs. Gene tree was built in RAxML and visualized in iTOL.

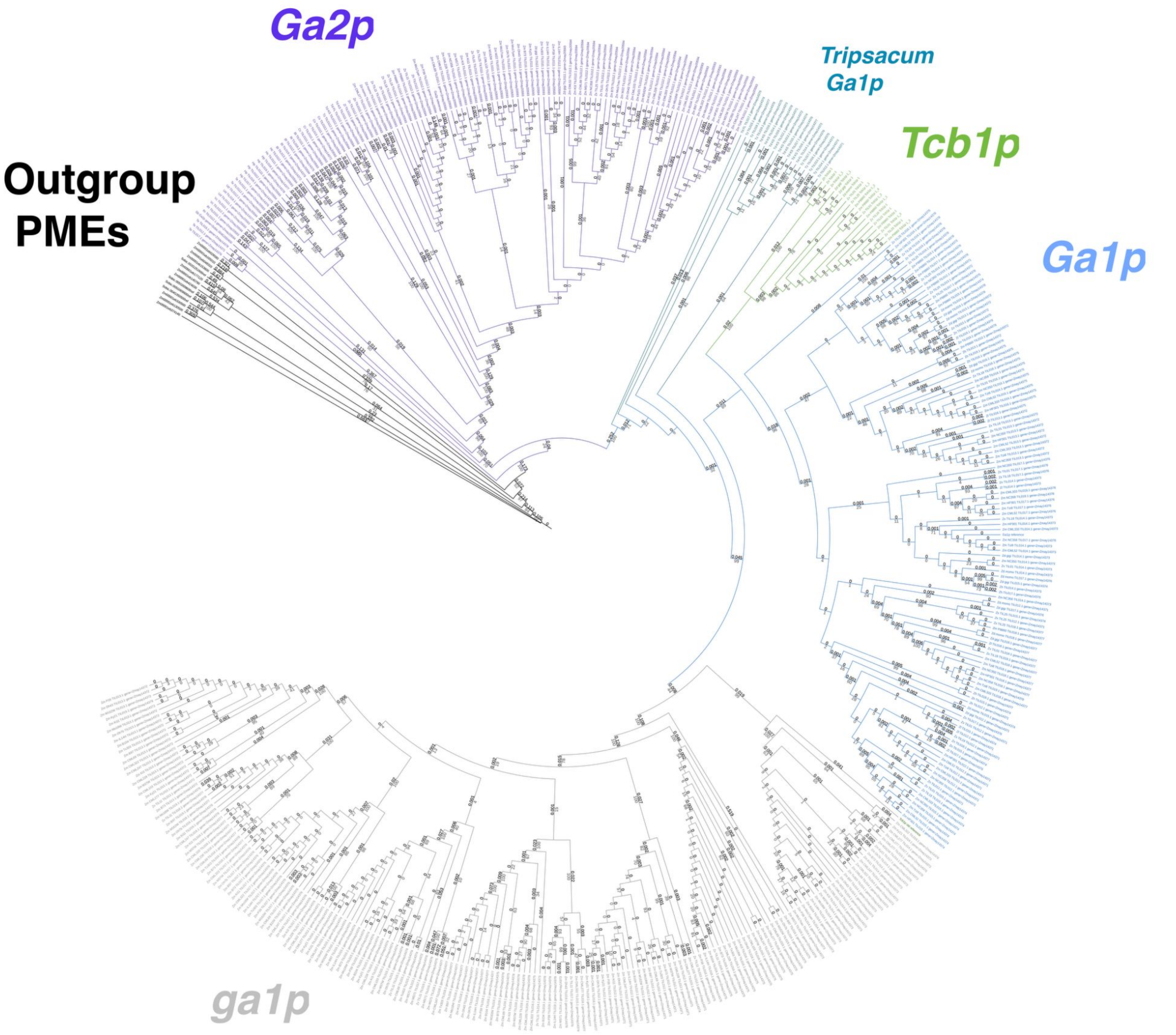


**Supplemental Figure 6b**
**GA pollen gene tree**
Gene tree of all full-length GA pollen gene sequences with *Zea mays mays* PMEs as an outgroup. Tree is based on nucleotide alignments of CDSs. Gene tree was built in RAxML and visualized in iTOL.

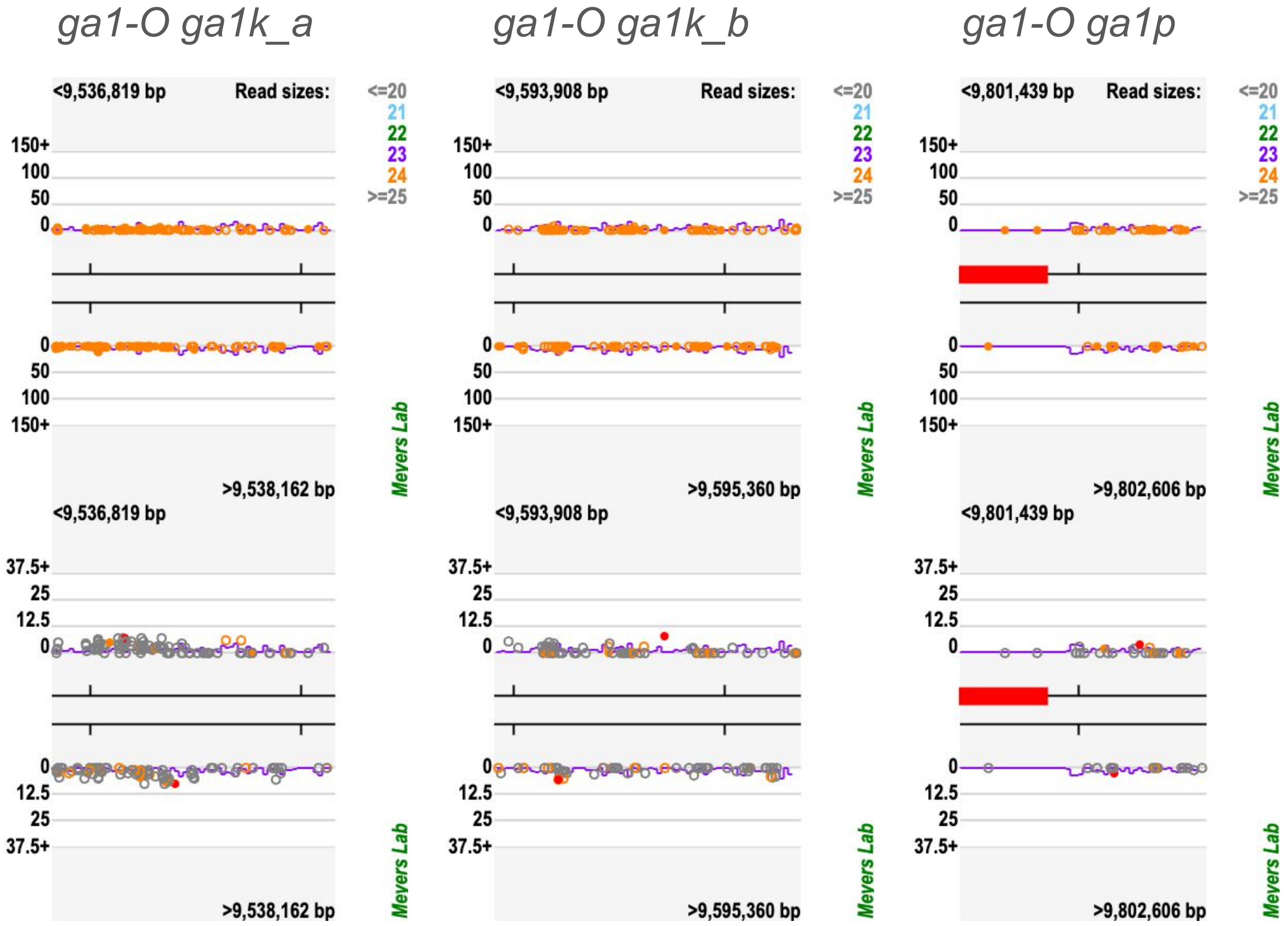


**Supplemental Figure 7**
**24nt siRNA hits and phase scoring for each ga1-O 'gene' in B73**

**24nt siRNA hits (upper panels):** Each dot represents a 24-nt siRNA hit, or regions in the B73 genome with exact sequence match to the siRNA. Hollow dots represent unique hits where the siRNA involved in the hit does not match the sequence of any other part of the B73 genome. Red bar on ga1p panel represents overlap with a B73 gene model for ZmPME3, which has been previously reported as the *Ga1* pollen factor. 24-nt siRNAs depicted are from the same lines and anther tissues described in main text. **Phasing Analysis (lower panels):** Each dot represents a "window" of ten cycles of small RNAs of length 24 nt, with the score for the degree of phasing indicated on the Y axis (scores calculated approximately as described by Howell et al., 2007). The red dot is the highest scoring window and has the best score in this region. Other colored dots are windows which are in phase with the highest scoring window -- exactly-in-phase windows appear as filled dots, and almost-in-phase (-1/+1) windows as hollow dots.

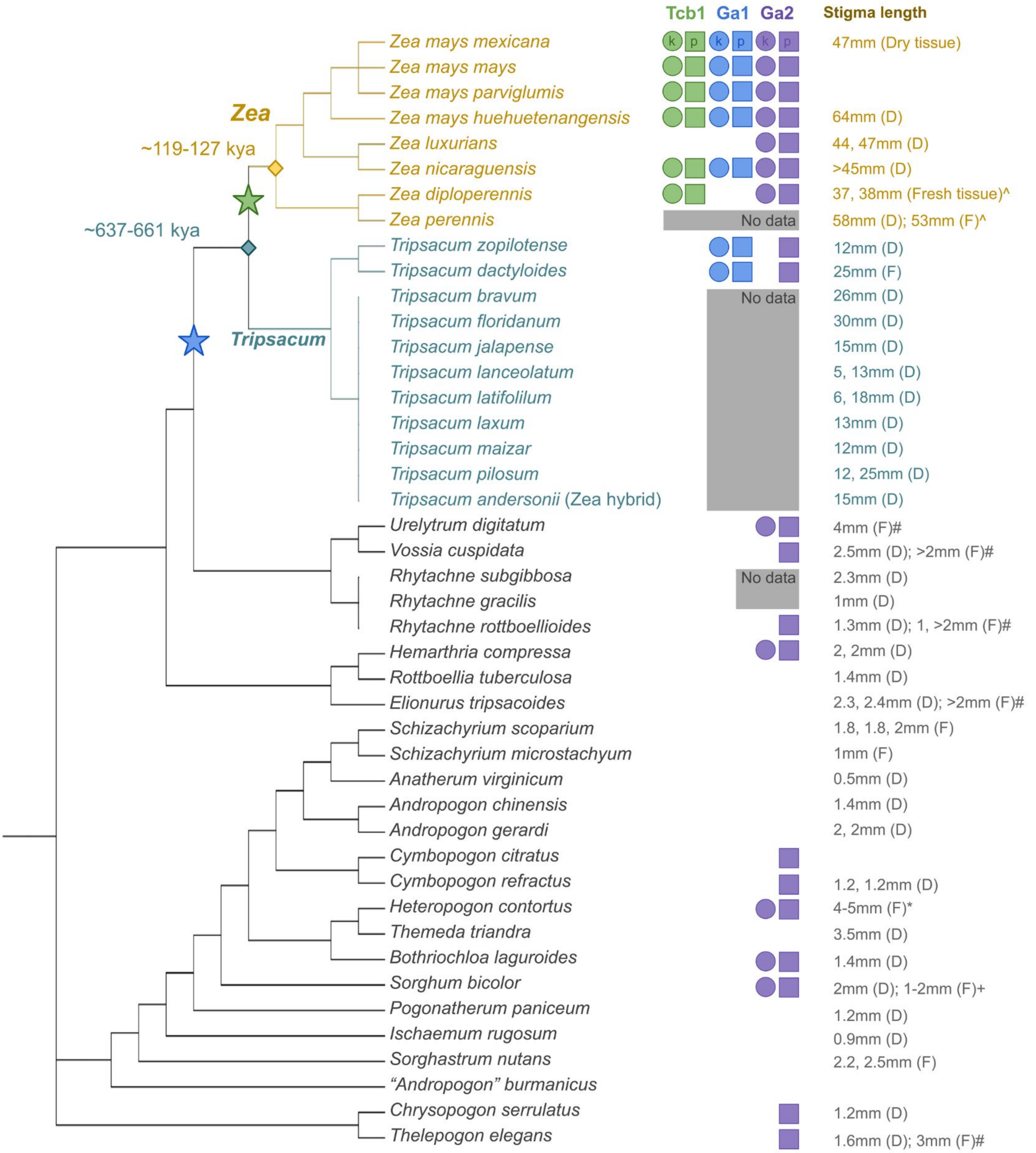


All stigma lengths measured by Elizabeth A Kellogg from fresh plant tissue or dried herbaria samples [see supplemental data]
Unless otherwise noted: * (Drisya and Pradeep 2020), + (Takanashi et al 2021),
### (Stapf 1917 Flora of Tropical Africa Vol IX Part 1 Ed. Sir David Prain), ^ (measured by Jeffrey Ross-Ibarra from fresh tissue)

#### Supplemental Figure 8

#### Stigma length and GA loci PAV on Andropogoneae species tree

Tree of all species we searched for GA loci gene sequences with stigma (silk) length and GA loci gene presence. For more information on GA PAV, species tree, and divergence times, see main text.

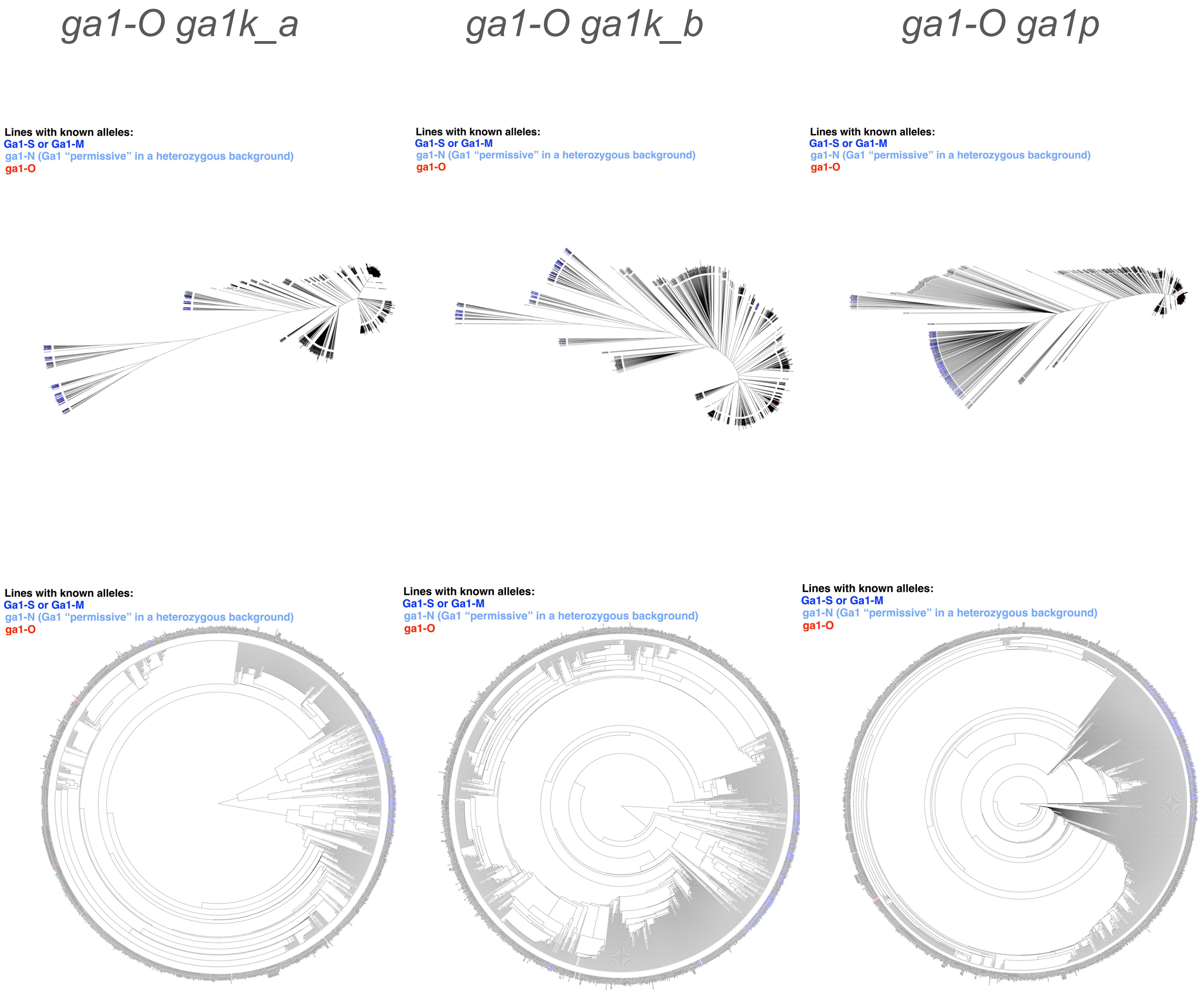


**Supplemental Figure 9**
**SNP trees for GA loci across diverse maize lines**
Neighbor-joining trees of SNPs in diverse maize lines alignment to the B73 *ga1k* and *ga1p* gene regions in the B73 *ga1-O* allele. Aligned SNPs were accessed through HapMap 4 on the Maize Genetics DataBase (MaizeGDB) website, which we also used to build the NJ trees. HapMap 4 and MaizeGDB:
Andorf et al 2024: https://doi.org/10.1101/2024.04.30.591904; Hufford et al 2021: doi:10.1126/science.abg5289; Grzybowski et al 2023: doi:10.1111/tpj.16123; Woodhouse et al 2021: doi:10.1186/s12870-021-03173-5
